# Sexually dimorphic neurons in the *Drosophila* whole-brain connectome

**DOI:** 10.1101/2025.06.10.658788

**Authors:** David Deutsch, Arie Matsliah, Anthony Crown, Kaiyu Wang, Sven Dorkenwald, Arpita Mondal, Subhajit Jana, Austin Burke, James Hebditch, Jay Gager, Edna Normand, Megan Wang, Szi-Chieh Yu, Amy Sterling, Claire McKellar, Philipp Schlegel, Stephan Gerhard, Gabriella Sterne, Marta Costa, Katharina Eichler, Yijie Yin, Gregory S.X.E. Jefferis, Barry Dickson, H. Sebastian Seung, Mala Murthy

## Abstract

Sexual dimorphisms in the brain arise from differences in cell number, morphology, and connectivity, and underlie sex-specific behaviors. In *Drosophila*, these differences are largely governed by the transcription factors *Fruitless* (Fru) and *Doublesex* (Dsx), expressed in neurons implicated in reproductive and social behaviors. However, how these neurons are integrated into the connectome to drive different behaviors in different sexes remains unclear. Here, we identify 98 putative Fru- or Dsx-expressing cell types (∼1700 neurons) in the female whole-brain connectome by matching electron microscopy reconstructions to light microscopy images. We then analyze their connectivity within the whole-brain network. Fru/Dsx neurons are strongly interconnected, but are not part of a private network. We measure distances to the sensory periphery to examine multisensory interactions, and map connections to descending neurons that drive behavior. We further cluster Fru/Dsx neurons based on whole-brain connectivity, and uncover functional groupings. We analyze a subset of strongly connected Fru/Dsx neurons to make functional predictions, and test some of these with experiments. Together, this work provides a brain-wide framework for understanding the organization of sexually dimorphic circuits and establishes a resource for dissecting the neural basis of social and sexual behaviors.

## Introduction

In *Drosophila melanogaster*, two transcription factors specify sex differences in brain development: Fruitless (Fru) and Doublesex (Dsx) ^1–6, 7^. Neurons that transcribe Fru and/or Dsx can be labeled in both males and females ^4,8–14^, revealing the population of neurons critical for sex-specific behaviors, such as courtship ^1,15–20^, receptivity ^8,21^, copulation ^22–24^, aggression ^4,25–29^, and egg laying ^20,30–33^ Fru/Dsx neurons have, by definition, sexually dimorphic gene expression. However, only a subset of the Fru/Dsx types are ‘sexually dimorphic’ in cell number, morphology, and connectivity ^34^. Previous work suggests that Fru and Dsx neurons preferentially connect to each other, though this has not been systematically evaluated. While targeted genetic reagents have enabled the study of specific cell types within this population as well as their connections to behavior, the recent release of a whole-brain connectome ^35,36^ makes it possible to analyze the connectivity of Fru and Dsx cell types within the larger network of the brain. As just one example, work on the Dsx+ vpoDNs, female-specific descending neurons that drive vaginal plate opening in sexually receptive flies, has revealed a circuit in which these neurons integrate courtship song information and mating-state signals to control receptivity behavior ^21^. But, how a cell type like vpoDN fits into the larger brain network (e.g., what other inputs to vpoDN are important for its activity) has not been systematically studied. Doing so relies on characterizing the connectivity of vpoDN (and other sex-specific or sexually dimorphic neurons) to all other neurons in the brain.

To identify putative Fru and Dsx neurons in a female whole-brain connectome ^36,37^, we compared light microscopic (LM) images ^10–12,21,30,36,38–42^ with electron microscopic (EM) reconstructions, a feat rendered possible by the stereotyped morphology of *Drosophila* neurons ^36^. We conducted this matching using tools for registering LM and EM datasets (see Methods; ^43,44^) and identified a population of 1574 Fru neurons, 50 Dsx neurons, and 78 neurons with morphology that fits both a Fru type and a Dsx type (denoted as Fru/Dsx). We map connectivity patterns of the Fru/Dsx neurons and show that they are more interconnected than spatially matched controls. We investigate how Fru/Dsx neurons combine sensory information across modalities and converge onto descending motor command neurons. We use connectivity to cluster these neurons and predict the function of new cell types. We then validate some of these predictions with experiments. We characterize both sex-specific neurons (present in only females) and sexually dimorphic neurons (present in both males and females) and provide a resource for future functional investigations of female sexual and social behaviors.

## Results

### Identifying Putative *Fruitless* or *Doublesex* neurons in the Female Whole-Brain Connectome

We first identified Fruitless (Fru) or Doublesex (Dsx) neurons (hereafter referred to as Fru/Dsx) by comparing light microscopy (LM) images of single or small subsets of Fru or Dsx neurons ^10–12,21,30,38–42,45,4610–12,21,30,38–41,45–47^ to electron microscopy (EM) reconstructions of single neurons in the connectome (Fig. 1A; see Methods and Table 1). When available, we also used previously labeled Fru/Dsx candidates in the ‘hemibrain’ dataset ^48^ as seeds for finding candidates in the whole-brain connectome (Supp Fig. 1 - S1). Additional Fru/Dsx candidates were then found in FlyWire using similarity in morphology or connectivity, as well as available annotations in Codex ^35,36,44^. In all cases, the final candidates were compared to LM images (see Table 1). Combining all available sources, we identified a total of 1702 putative Fru/Dsx cells in the whole-brain connectome belonging to 6 super classes (Figs. 1B-C and Fig. 2). These neurons make up 98 Fru/Dsx types (Fig. 2), with roughly the same number of cells per hemisphere for each type (Fig. 1D), and cell bodies present in both the anterior and posterior female brain (Fig. 1E-G). In many cases, a given LM cell type (by morphology) corresponds to multiple possible cell types in the connectome (a connectome cell type is largely defined by connectivity ^36,49^), resulting in multiple ‘subtypes’ for a single Fru/Dsx type and a total of 272 subtypes (Fig. 1A, Supp Fig. 1 - S1-3 and Table 1). We include all EM candidates with LM matches that exhibit strong anatomical similarities to the LM images (Supp Fig. 2 - S1-2), generally preferring false negatives over false positives (e.g., Supp Fig. 1 - S6, Supp Fig. 2 - S3). In two cases, Fru and Dsx types have shared morphology and we group them together in our analyses: Dsx pCd1 neurons share morphology with Fru pMP3, and Dsx pCd2 neurons share morphology with Fru pMP5 (Supp Fig. 1 - S2). All of our labels have been available in Codex (codex.flywire.ai) since the release of the connectome dataset ^35,36^. As our identification is based solely on morphological matching between LM and EM, it is possible that some putative Fru/Dsx neurons in our list do not actually express the Fru or Dsx transcripts (see Discussion).

**Figure 1.**
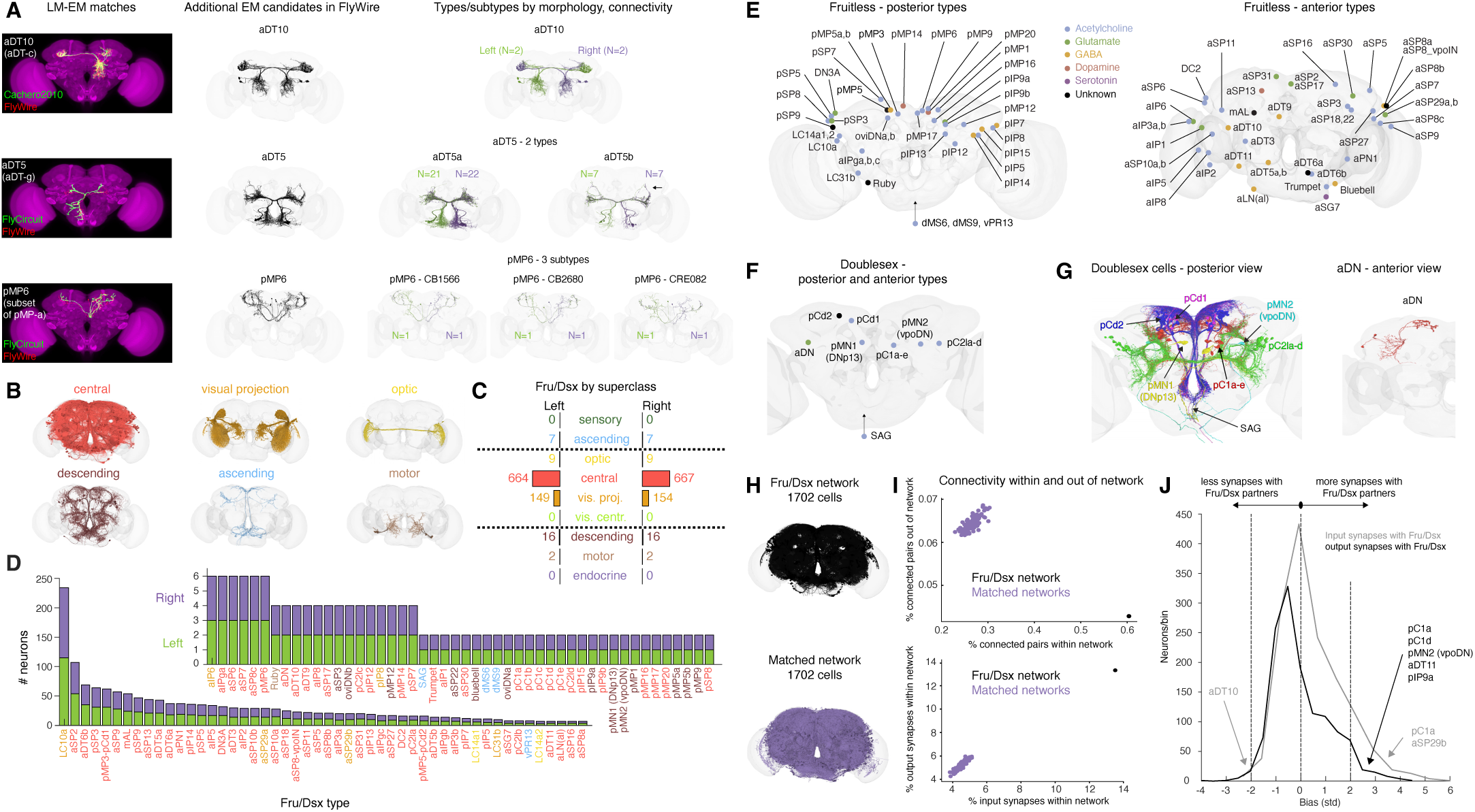
Identifying putative *Fruitless* and *Doublesex* (Fru/Dsx) neurons in the FlyWire brain connectome. **(A)** FlyWire (EM) candidates of Fru/Dsx types were found by direct comparison to light microscopy (LM) stacks of Fru/Dsx neurons (single neuron ‘clones’ or sparse neuron labeling with genetic lines ^10–12,21,30,38–41,45,52^) or via matching to previously identified Fru/Dsx candidates in the hemibrain dataset ^36,48^. In both cases, FlyWire candidates were compared with all available sources (see Table 1). Additional Fru/Dsx candidate neurons in FlyWire were identified based on existing annotations of FlyWire neurons in Codex ^36,49^; see Methods). For a given LM source there may by a single Fru/Dsx type (e.g., aDT10; top row), or multiple types that both overlap with the LM morphology, but are not similar in morphology (e.g., Fru/Dsx aDT5a and aDT5b; middle row). Since Fru/Dsx types are defined only by morphology, a given Fru/Dsx may comprise multiple FlyWire subtypes that differ by their connectivity (e.g., pMP6; bottom row). There are a total of 98 Fru/Dsx types and 272 subtypes in the dataset. **(B)** We identified 1702 putative Fru/Dsx cells in total, here sorted by ‘superclass’ ^35^ and rendered to a brain template. Our dataset excludes Fru/Dsx neurons in the mushroom body (see Methods), and in the following superclasses: sensory and visual centrifugal (see codex.flywire.ai for more information on superclasses). (**C)** Number of Fru/Dsx cells in each superclass and hemisphere. **(D)** Number of cells per type in the left (green) and right (purple) hemispheres. Type names are colored by superclass (as in B-C). **(E)** Soma clusters of putative Fru types in the posterior (left) and anterior (right) brain, colored by known or predicted ^53^ neurotransmitter expressions. When classifier prediction did not agree on at least 60% of individual neurons, the type was colored back. **(F)** same as in (E), but for all Dsx types, in posterior view (all Dsx cells in the brain are posterior, except aDN). The Fru ascending dMS6, dMS9 and vPR13 and the Dsx ascending SAG neurons are marked outside the brain (not at their physical location) with upward arrows. Fru pMP3 and Dsx pCd1 share morphology, as do Fru pMP5 and Dsx pCd2. Because we are unable to distinguish these types in the connectome, they are referred to as pMP3/pCd1 and pMP5/pCd2. While pMP5/pCd2 has 16 cells, there are thought to be only 4 Dsx pCd2 neurons^46,54^ (note that Dsx pCd neurons are also termed ‘pC3’^5,19^). Dsx pCd2 neurons are downstream of Dsx pC1a neurons and upstream of Fru Lgr3-expressing cells^50,55^ which are morphologically similar to the aDT6a,b neurons (see also ^7^). Only 2 pMP5/pCd2 subtypes (SMP286 and CB0405; 4 cells in total) are directly downstream of pC1a and upstream of aDT6a,b, making them good candidates for being Dsx pCd2 cells. Those subtypes also have over 50% of both their inputs and outputs with Fru or Dsx cells, compared to 10% or less for some other pMP5/pCd2 subtypes.(**G**) All posterior (left) and anterior (right) putative Dsx neurons rendered to a brain template and colored by type. aDN neurons (2 per hemisphere) are shown only in the left hemisphere. Since we couldn’t distinguish morphologically between Dsx pCd1 and Fru pMP3 or between Dsx pCd2 and Fru pMP5 (Supp Fig. 1 - S2), they are depicted separately in the Fru and Dsx soma-location maps (Fig. 1E-G). The Dsx types pMN1/pMN2 ^17,46^ have identical morphology to the DNp13/vpoDN types ^21,23,56^, respectively. **(H)** All 1702 putative Fru/Dsx neurons (black), and a set of 1702 neurons from one example ‘matched network’ (purple). ‘Matched networks’ (see Methods) have the same superclass and spatial distribution as the Fru/Dsx neurons. **(I)** (top) Fru/Dsx neurons (black) make more connections within-network (versus out-of-network), as compared with 100 matched networks. A 5 synapse minimum threshold for a connection was applied. (bottom) Fru/Dsx neurons (black) have more input and output synapses within-network, as compared with 100 matched networks (purple). Synapses within a primary type were excluded for both Fru/Dsx and matched networks, to avoid possible bias for within network connectivity due to more cells per type in the Fru/Dsx network compared to a matched network. **(J)** Synaptic Input (gray) and output (black) bias distributions, measuring the bias of each Fru/Dsx type to form more (positive bias) or less (negative bias) synapses with Fru/Dsx partners compared to the number of synapses with any of its partners (see Methods). The input and output biases were calculated separately for each individual Fru/Dsx cell, and then averaged for all the cells of a given type. For example, pC1a ^21^ has a strong bias for connections with Fru/Dsx partners for both inputs and outputs, while for pC1d, the strong bias is only for the output connections ^25,27^. aDT10 has a negative input bias.

**Figure 2.**
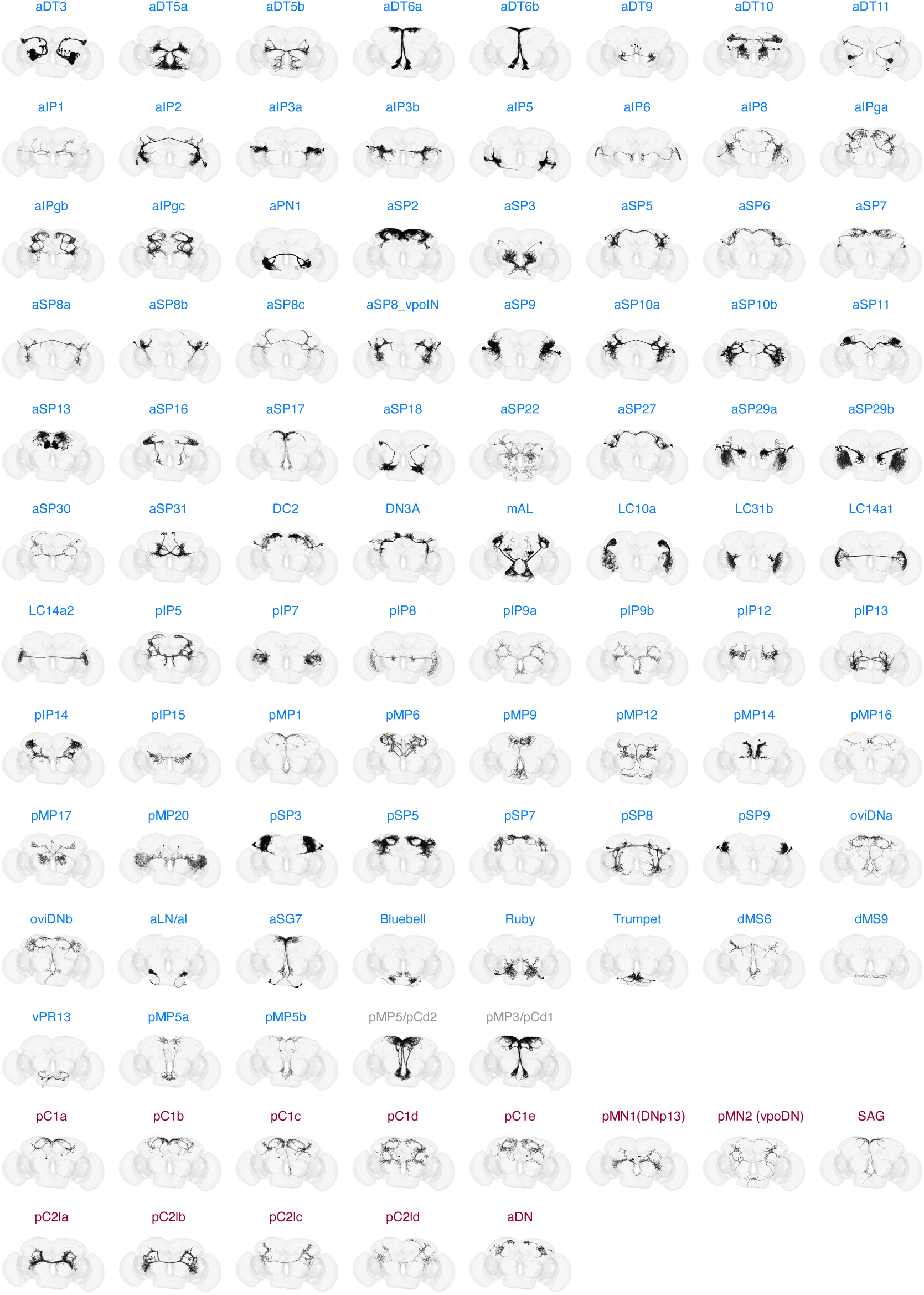
EM renders of Fru/Dsx types. All Fru/Dsx types rendered in FlyWire. All the neurons for a given type are shown. Fru/Dsx type names are colored blue, red, and gray for ‘Fru’, ‘Dsx’ and ‘Fru or Dsx’ as elsewhere in the manuscript. For type LC10a, only a subset is shown.

### Connectivity Within the Fru/Dsx Network

Prior work on individual Fru or Dsx cell types has focused mostly on connections with other Fru/Dsx neurons ^16,21,25,27,50,51^. This focus has led to an assumption that Fru/Dsx neurons preferentially connect to each other ^11^. We tested this assumption first by building ‘matched networks’: random sets of neurons in the whole-brain connectome that match the spatial and super class distributions of Fru/Dsx neurons and with the same total number of cells (Fig. 1H and Supp Fig. 1 - S4; see Methods). To compare these networks, we constructed two metrics for their interconnectivity. First, we measured the fraction of synapses between each cell and other cells in its network (Fig. 1I, top). Second, we compared the fraction of connected cell pairs in each network out of all possible pairs (Fig. 1I, bottom). Both measures indicate that Fru/Dsx cells are significantly more interconnected than expected based on their physical proximity. We uncovered a strong positive bias for both inputs and outputs of Fru/Dsx neurons (Fig. 1J and Supp Fig. 1 - S5) - that is, most Fru/Dsx neurons make stronger connections with their Fru/Dsx partners than with non-Fru/Dsx neurons. There is heterogeneity, however; some cell types have a very strong positive bias for both their inputs and outputs, some have a strong positive bias for only their inputs or outputs, and a single type (aDT10) has a negative input bias. The heterogeneity of the Fru/Dsx network can be appreciated by examining the fraction of input and output synapses with other Fru/Dsx neurons (Supp Fig. 1 – S3). Some types, like SAG or pC1a, have mostly connections with other Fru/Dsx neurons, while other types, like pSP8 or aIP6, do not; notably SAG and pC1a neurons are both Dsx+ and we show below that Dsx+ cell types are over-represented in hubs in the network (Fig. 6). This analysis may provide information for sorting which subtypes are more likely to express Fru or Dsx (Supp Fig. 1 – S3A), if we assume that Fru/Dsx neurons are more likely to connect with other Fru/Dsx neurons. We also identified neurons in the connectome that are not labeled as Fru/Dsx, but that make a large percentage of their synapses with Fru/Dsx neurons (Supp Fig. 1 – S3B); this list may contain false negatives.

While the total number of Fru/Dsx neurons in the female brain is not known, prior work estimates 1500-3000 neurons ^10,11,57^ or roughly 2% ^58^ of the neurons in the female brain are Fru while a few tens are Dsx ^19,46^. Based on these estimations, we have likely identified just over 50% of the Fru/Dsx neurons in the female connectome. This underrepresentation is due to i) exclusion of both sensory neurons and mushroom body Kenyon cells, as we could not unambiguously identify the Fru subsets among them (similar to what was done in ^11^), ii) exclusion of many optic lobe neurons that were similarly difficult to distinguish Fru from non-Fru ^49,59^ (e.g., the medulla ‘M neuron’ ^39,60^), and iii) exclusion of putative Fru/Dsx types for which we did not have LM images. Because the FlyWire brain connectome does not contain information beyond the neck connective, a full CNS connectome is necessary to identify more Fru/Dsx ANs with cell bodies and dendrites in the nerve cord ^42^. Many Fru/Dsx cell types were named previously ^3,5,10–12,21^ and we used these existing names where possible (see Table 1), primarily using the nomenclature from Yu et al. (2010) ^11,12^, and assigning new names following this convention. For example, aSP30 is a previously uncharacterized group (see Fig. 7 for proposed function), but its cell bodies are in the vicinity of other aSP Fru types, such as aSP3 ^11^ and aSP18 ^12^. We identified 85 Fru cell types (Fig. 1E and Fig. 2) and 15 Dsx cell types, including the ascending SAG neurons (Figs. 1F-G and Fig. 2). The division of pC1 cells into subtypes was previously reported ^21,25^, while the division of pC2l is newly described in this study (see Fig. 7). We generated a comprehensive online catalog (https://codex.flywire.ai/research/dsx_fru?dataset=fafb) with an ‘ID card’ for each type that contains information about cell morphology, known or predicted neurotransmitter expression ^53^, and top inputs and outputs in the whole-brain connectome (from any cell type).

### Sensory Processing in the Fru/Dsx Network

Prior work used a probabilistic model to estimate information flow in the connectome, from sensory populations through to different intrinsic neurons and out to motor or descending neurons (Supp Fig. 3 - S1)^35^. This method generated a mapping of every neuron in the central brain (not including the optic lobes) relative to the sensory periphery (or sets of sensory seed neurons)^35^; Fru/Dsx neurons are distributed throughout this map (Fig. 3A, Supp Fig. 3 - S2). Here we inspect this ordering to make predictions about the responsiveness of Fru/Dsx types to different sensory modalities (see comments on the interpretation of rank analysis under Discussion). By convention, rank 1 cells are defined as cells inside the seed (sensory) group. While it is known that some sensory neurons express Fru^2,11,15,58,61–64^, as stated earlier, we did not include these neurons in our Fru/Dsx list.

**Figure 3.**
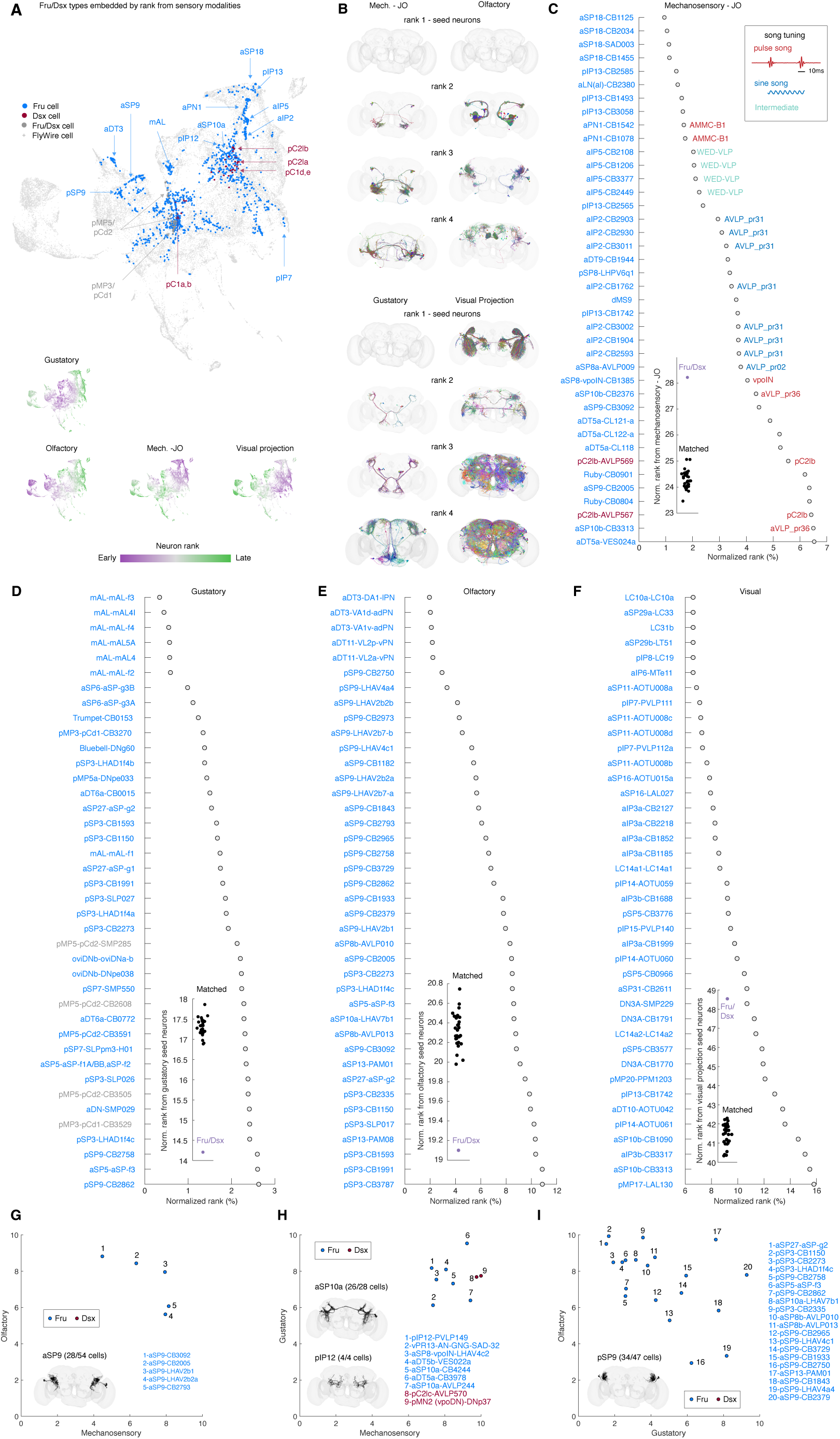
Information flow from sensory modalities to Fru/Dsx neurons. **(A)** Top, UMAP analysis of the matrix of normalized traversal distances ^35,80^, resulting in a 2D representation of each Fru/Dsx cell excluding visual projection and ascending neurons in a sensory space as in Dorkenwald et al. 2024^35^ (see Methods). Each dot represents a single neuron. Fru, Dsx, Fru/Dsx neurons are colored in blue, red and gray, respectively, while any other FlyWire cell is represented as a gray cross. The approximated average location of the cells belonging to some types are shown. The Bottom, we coloured neurons in the UMAP plot by the rank order in which they are reached from 4 seed groups (Mech. -JO: mechanosensory-JO; see ^35^). Purple neurons are reached earlier than green neurons. For example, mAL neurons (see arrow in top panel) are positioned around cells with early rank from the Gustatory sensory neurons (bottom panel ‘Gustatory’), consistent with their role in processing gustatory cues ^15^, while aDT3 has small rank from the olfactory seed ^11,63,70^, while aSP18 has small rank from the Mech. - JO seed, consistent with its low rank from Mechanosensory-JO (see 7(C); ^12^). **(B)** Fru/Dsx neurons with different ranks from different sensory seeds (mechanosensory-JO, olfactory, gustatory, or visual projection; see Methods). No mechanosensory-JO, olfactory and gustatory sensory neurons (rank 1) exist in our Fru/Dsx network, while some visual projection neurons do. **(C)** Fru/Dsx subtypes are sorted by normalized rank from Mechanosensory-JO. Known auditory types ^17,67^ are colored according to their tuning to syllables of courtship song, ‘pulse’ or ‘sine’. Intermediate turning implies no strong preference to Pulse over Sine song as previously defined ^67^. **(D-F)** same as (C) for Gustatory (D), Olfactory (E), and Visual projection (F) seeds. Fru/Dsx subtypes are colored as elsewhere. Normalized ranks from sensory modalities were calculated as before (see Methods), by converting absolute ranks to percentile as previously done ^35^, and averaged over all the cells in a given subtype. For example, aSP18-CB1125 has a normalized rank of ∼1% from mechanosensory-JO, implying that ∼99% of FlyWire cells have a higher absolute rank than an average cell in the aSP18-CB1125 group. **(G–I)** Fru/Dsx subtypes with normalized rank < 10% with pairs of modalities: mechanosensory-JO and olfactory (G), mechanosensory-JO and gustatory (H), gustatory and olfactory (I). Insets show rendered neurons for some multisensory types. Only around half of the aSP9 neurons (28/54) are multisensory (according to their ranks) compared to 26/28 for aSP10a and 4/4 cells for pIP12. Pairs with visual-projection are not shown: no type has normalized rank <10% from both visual proj. and gustatory seeds. Types pSP5-CB3776 and aIP3b-CB1688 have norm. rank<10% with vis. Proj. and norm. rank <10% with olfactory or mech.-JO seeds, respectively.

We seeded Fru/Dsx subtypes from 4 sensory modalities (Fig. 3B-F): mechanosensory (in which we use as our seed the Johnston’s Organ neurons (JONs) of the antenna ^65,66^), gustatory and olfactory (using gustatory and olfactory sensory neurons as seeds), and visual (in which we use all visual projection neurons as the seed ^35^). Some Fru/Dsx cell types that are close to the sensory periphery have been previously studied and shown to respond to particular sensory stimuli, including Fru aLN(al), aPN1 (a subset of AMMC-B1), and aIP5 (also annotated as WED-VLP ^67^) neurons in the auditory pathway ^67–69^ and aDT3 and aDT11 (also called VL2a_vPN/VL2p_vPN) in the olfactory pathway ^58, 70,71^. In the gustatory system, mAL neurons have been studied in males for their role in driving arousal following tasting/tapping of the female ^15,72^ but their role in females is uncharacterized. In the visual system, Fru LC10a and LC31b are visual projection neurons (VPNs). While the role of LC10a was characterized ^73^, the role of LC31b is unknown (but see Fig. 7). Rank distance to each sensory modality serves as a resource for making functional predictions. For example, we predict aSP18 and pIP13 are mechanosensory, while aSP6 is gustatory, and aSP11, pIP7 (postsynaptic to LPLC2 ^74^), and aIP3a (postsynaptic to LC16^75^) are all visual.

The small distance of oviDN neurons^30^ from the taste periphery suggests a pathway by which taste information influences egg laying decisions. Fru/Dsx neurons have lower rank from the olfactory and gustatory seeds compared to matched networks (Fig. 3D-E), and higher distance from the visual and mechanosensory seeds (Fig. 3C and F). It is important to note that rank distance is a probabilistic measure, and cells with high rank from a given modality may still be weakly connected directly to ‘seed’ neurons of this modality. For example, aSP9-LHAV2b2a receives weak direct input from LC43, but is of high normalized rank from the visual projection neurons (Fig. 3F).

Social behaviors in *Drosophila* often involve integration of sensory cues of different modalities ^2,8,15,76^. By examining the proximity of the Fru/Dsx neurons to more than one ‘seed’ modality (Fig. 3G-I, Supp Fig. 3 - S3), we find Fru/Dsx neurons that are close to both olfactory and mechanosensory receptor neurons (Fig. 3G), to both gustatory and mechanosensory receptor neurons (Fig. 3H), and to both olfactory and gustatory receptor neurons (Fig. 3I). There were no visual-gustatory neurons (with ranks <10%, see Methods), only 1 visual-olfactory cell type and 1 visual-mechanosensory cell type (see Figure legend). aSP9 neurons are predicted to process both olfactory and mechanosensory information: they primarily receive inputs in the lateral horn (LH; containing third order olfactory neurons) and project to the AVLP and PVLP; all three brain areas are known to carry auditory signals ^77^, and the LH is a major area for olfactory processing ^74^. While prior work showed that vpoIN, vpoDN, and aSPp10a respond to courtship song (auditory) stimuli ^21,67^, here we predict these neurons should also process gustatory information. Further, we predict that pSP9 neurons in the lateral horn process both olfactory and gustatory information, but only the olfactory coding properties of these neurons have been studied ^70,78,79^.

### Connectivity With Descending Neurons

In *Drosophila* and other insects, the neck connective is both a physical and informational bottleneck connecting the brain and the ventral nerve cord (VNC; spinal cord analogue) ^42,81–83^. While some descending neurons (DNs; carrying information from the brain to the VNC) and ascending neurons (ANs; carrying information from the VNC to the brain) have been previously identified as Fru/Dsx (refs), we identify additional neurons here. In total, we label 14 ascending (4 types) and 32 descending (13 types) as Fru/Dsx cells. We then mapped direct connectivity between all Fru/Dsx subtypes (Figs. 1-2) and any descending neuron in the whole-brain connectome, focusing on the strongest connections (Fig. 4; see Methods), in order to identify DNs that may be important for sexual and social behaviors. We find that 31 DN types have 10% or more of their input synapses with Fru/Dsx neurons (including Fru/Dsx DNs). Examining the network of DNs connected to Fru/Dsx neurons reveals several interesting features. First, while we do not yet know the function of DNpe044, and DNpe046, their clustering with DNs involved in oviposition (pMP1, oviDNa, and oviDNb) suggests they are involved in this behavior. DNp68, similarly, has never been studied, but receives input from pC1d and aIPg neurons, previously studied for their role in female aggression ^25–27^, as well as pC2l, important for song responses^17^. We find that DNp13 neurons receive inputs that are similar to DNpe050 and DNpe052, two DN populations that have not yet been characterized; they may similarly be involved in rejection in response to audiovisual cues ^56^. We find that pC1a, whose activation drives the vpoDNs which mediate sexual receptivity to male song ^21^, also drives pMP12-DNp60, implicating this new cell type in sexual receptivity.

**Figure 4.**
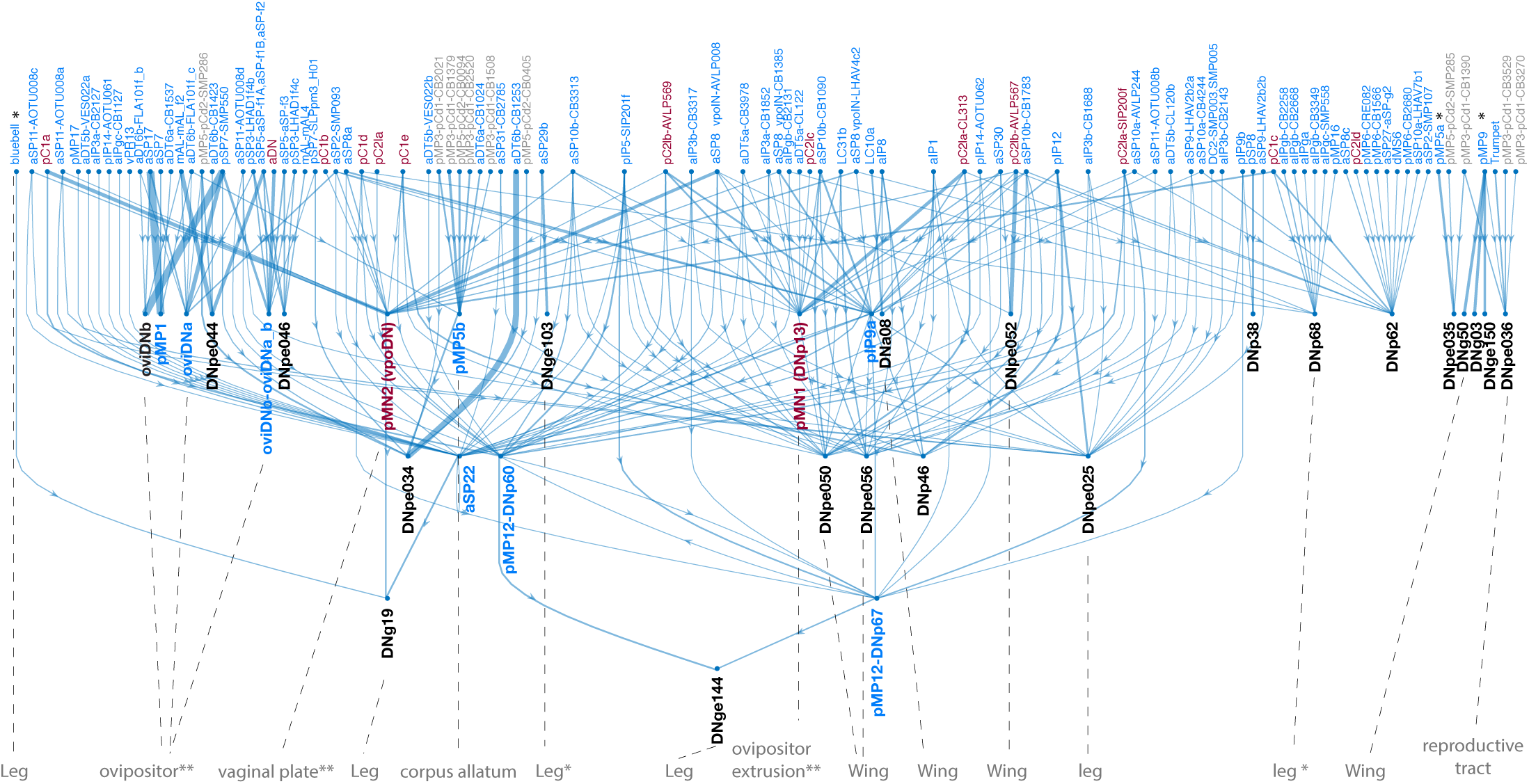
Fru/Dsx neuron connectivity to descending neurons. Strong direct synaptic connections (1% or larger, symmetric threshold; see Methods) between Fru/Dsx neurons and DNs (including DNs that are Fru/Dsx, such as pMN1-DNp13 and pMN2-vpoDN and DNs that are not in the Fru/Dsx list). This map reveals DNs that may be important for driving sexually dimorphic behaviors. Some of these have already been characterized (for example, Fru oviDNs or Dsx vpoDN ^21,30,31^), but most have not. Fru, Dsx, Fru/Dsx types are colored blue, brown and gray as elsewhere. Descending neurons not in the Fru/Dsx list are marked in black. Possibly some of these types are Fru or Dsx positive. All the cell types that are not in the top row of the map are descending. Descending types that are in the top row are marked with (*). As some descending neurons are upstream of other descending neurons, they are not at the bottom row of the map. One Fru/Dsx descending type is missing from the diagram: aSP3 (DNa13); this is because it does not match the threshold of 10% inputs from Fru/Dsx neurons to a DN applied for this Figure. Predicted effector (bottom line) is based on BANC ^84^ v626 (through CODEX). Predicted effector is not shown when not consistent across cells of the same descending type (e.g., for DNg03 ‘middle_leg_neurosecretory_cell’ is predicted for one cell and ’wing_neurosecretory_cell’ is predicted for the other). * similar but not identical effector is predicted (e.g., for DNge144 the prediction for one cell is ‘middle_leg_motor’ and for the other ‘front leg motor’’). ** prediction made based on literature (e.g., DNp13 controls ovipositor extrusion while vpoDN controls vaginal plate opening).

Finally, we identified aSP30 in this study as a cross-pathway inhibitory neuron (Fig. 7) - it receives excitatory input from neurons involved in receptivity and inhibits neurons involved in rejection (like DNp13). And, we find that aSP30 also inhibits both DNpe050 and DNpe052, further implicating these descending neurons in rejection behaviors. Lastly, though not specific to Fru/Dsx connectivity, is that several DNs have other DNs as one of their major inputs (Supp Fig. 4 - S1). This type of DN-to-DN connectivity has been recently analyzed functionally ^83^ and is likely important for recruitment of motor command pools to drive behaviors. The recent completion of a full CNS (brain and nerve cord) connectome ^84^ renders it possible to interpret the connections shown in Figure 4 by analyzing the outputs of these DNs onto the motor circuits of the VNC (Fig. 4).

A total of 17 sexually dimorphic or female specific DN types with identified projections in FAFB ^85^/FlyWire ^35,36^ were recently reported based on the comparison of male and female nerve cord connectomes ^42^. Most of those are included in our Fru/Dsx list. Of those not included in our Fru/Dsx list, some are shown to be strongly connected to the Fru/Dsx network (e.g., DNpe044, DNpe046; Fig. 4A). Future studies will reveal if those neurons express the fruitless gene.

### Connectivity-Based Clustering of Fru/Dsx Neurons

Previous work on the optic lobe has revealed that the connectivity of all neurons can be used to cluster them into functional groupings ^49^. We applied the same method to cluster the Fru/Dsx network based on their connectivity to all neurons in the brain connectome (Supp Fig. 5 - S1). Using this method, 272 Fru/Dsx subtypes can be sorted into 27 groups (Fig. 5). Our method groups similar subtypes, as expected. For example, the aSP9 neurons consist of 12 subtypes, together composing a single cluster (cluster 7), while the subtypes that compose the aIPa,b,c neurons are all in cluster 10. Further, aIPg neurons are clustered with the LC10a neurons, both shown to be involved in aggressive pursuit ^25,27,86^. Most of the Dsx types are contained in two clusters (clusters 1 and 16). Cluster 1 contains the SAG neurons, pC1a/c, and pMN2/vpoDN which are all involved in the control of female receptivity ^21,87^. Cluster 16 includes the types pC1d/e, pC2l, and pMN1/DNp13, which are involved in rejection/aggressive behaviors and also shown to respond to courtship pulse song^17,23,25,27,56^ (male flies sing song sequences consisting of switches between sine and pulse song ^88^). We find that clusters also sort by sensory modality. Clusters 12 and 24-25 contain auditory neurons (e.g., aLN/al ^68,69^), clusters 5 and 22 contain gustatory and visual neurons (e.g. mAL ^15^ and LC14 ^40^), respectively, while clusters 6-8 contain olfactory neurons (e.g., aDT3-DA1 ^89^). Some subtypes are split between clusters, indicating heterogeneity. For example, cells that are either pMP3-pCd1 (excitatory) or pMP5-pCd2 (inhibitory) separate into two clusters and are also heterogeneous in their connectivity with other Fru/Dsx neurons (Supp Fig. 1 - S3). The 11 pMP3-pCd1 and pMP5-pCd2 neurons that are most strongly connected with other Fru/Dsx neurons are all in Cluster 1, which also contains multiple Dsx neurons. Meanwhile, cluster 3 contains the pMP3-pCd1 and pMP5-pCd2 cells that are less connected to other Fru/Dsx neurons. This pattern of segregation among the clusters may indicate which subtypes of pMP3-pCd1 and pMP5-pCd2 are more likely to be Dsx+ versus Fru+.

**Figure 5.**
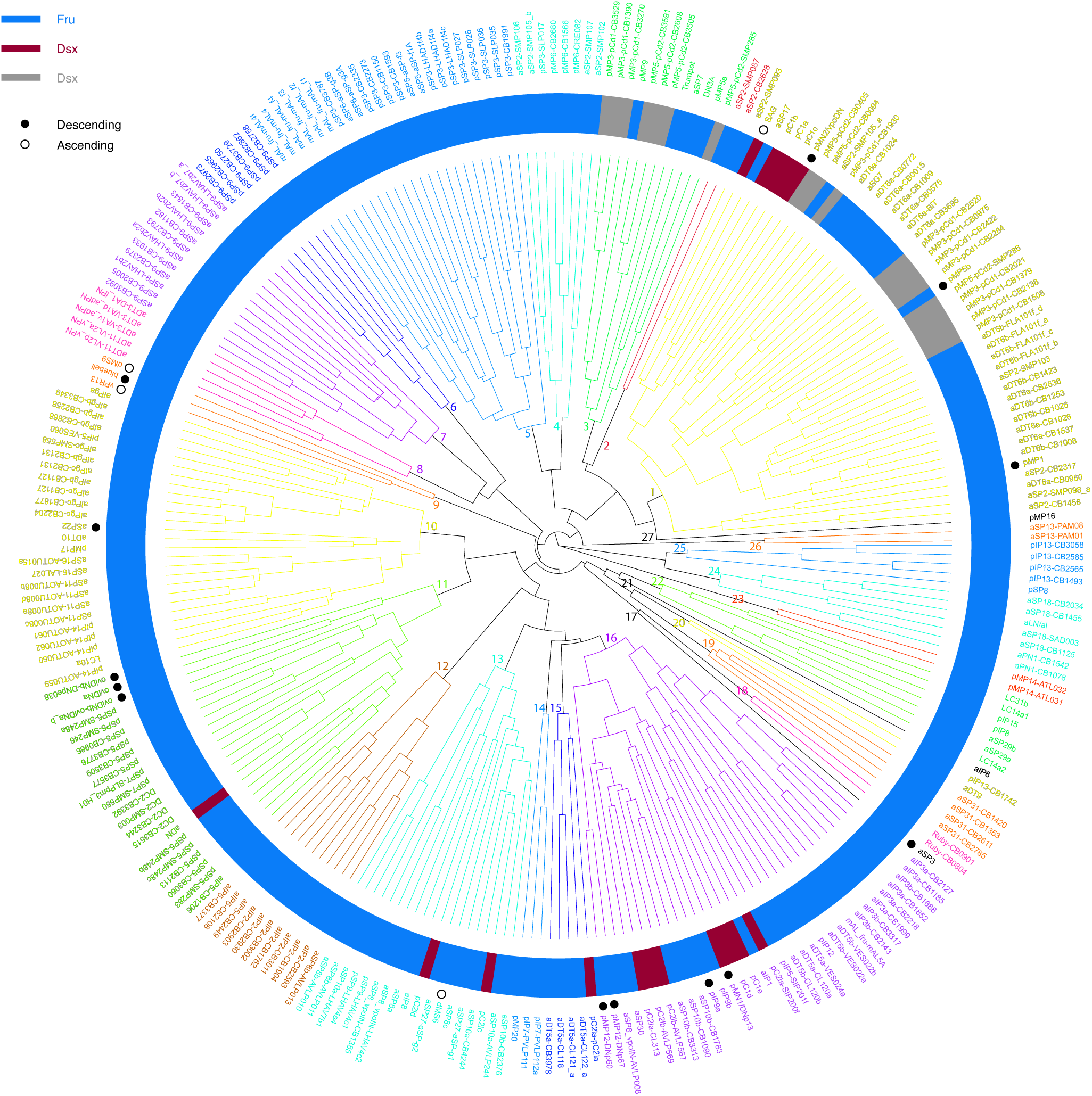
Connectivity-based clustering of Fru/Dsx neurons. Fru/Dsx neurons were clustered into 27 clusters based on their connectivity (input and output) with all cell types in the whole brain connectome (see Methods). Cell type names are colored by cluster. Fru cell types are indicated in the inner ring with blue rectangles, Dsx cell types with dark red rectangles, and Fru/Dsx cell types with gray rectangles. Consistent with previous clustering into subtypes ^36,44^, cells from the same primary type (or Fru/Dsx subtype) typically share the same cluster. A black filled circle indicates a descending type, a black empty circle indicates an ascending type.

Next, we use the clusters to construct a simplified network diagram (Fig. 6) by considering each one of the 27 clusters as a single node (see Methods). In short, weights are calculated based on normalized synapse numbers and nodes are positioned such that backward connections are minimized. While nodes are positioned based on all weights, only the strongest ones are shown. Therefore weakly connected nodes (nodes 18, 20, 21, 23) are shown as disconnected. Based on known literature and on rank distance from sensory modalities (Fig. 3) we infer the role of some nodes. For example, cluster 17 includes a single type, the aSP3 neurons (4 descending neurons also called DNa13 ^42^) that target the three leg neuropils ^90^ and may be key in controlling female walking in response to visual cues (connection to cluster 22) in the context of rejection/aggressive behaviors (direct projections from clusters 10,16). The first layer of the simplified network contains neurons that have small rank distance (see ^35,80^) from olfactory, gustatory, and auditory sensory neurons. Node 13 is multisensory, containing both known auditory neurons (pC2l, aSP10 ^67^) and Fru types with small rank distance from mechanosensory and olfactory sensory neurons (pSP9-LHAV4c1, pSP9-LHAV4a4 ^91^; see Fig. 3).

**Figure 6.**
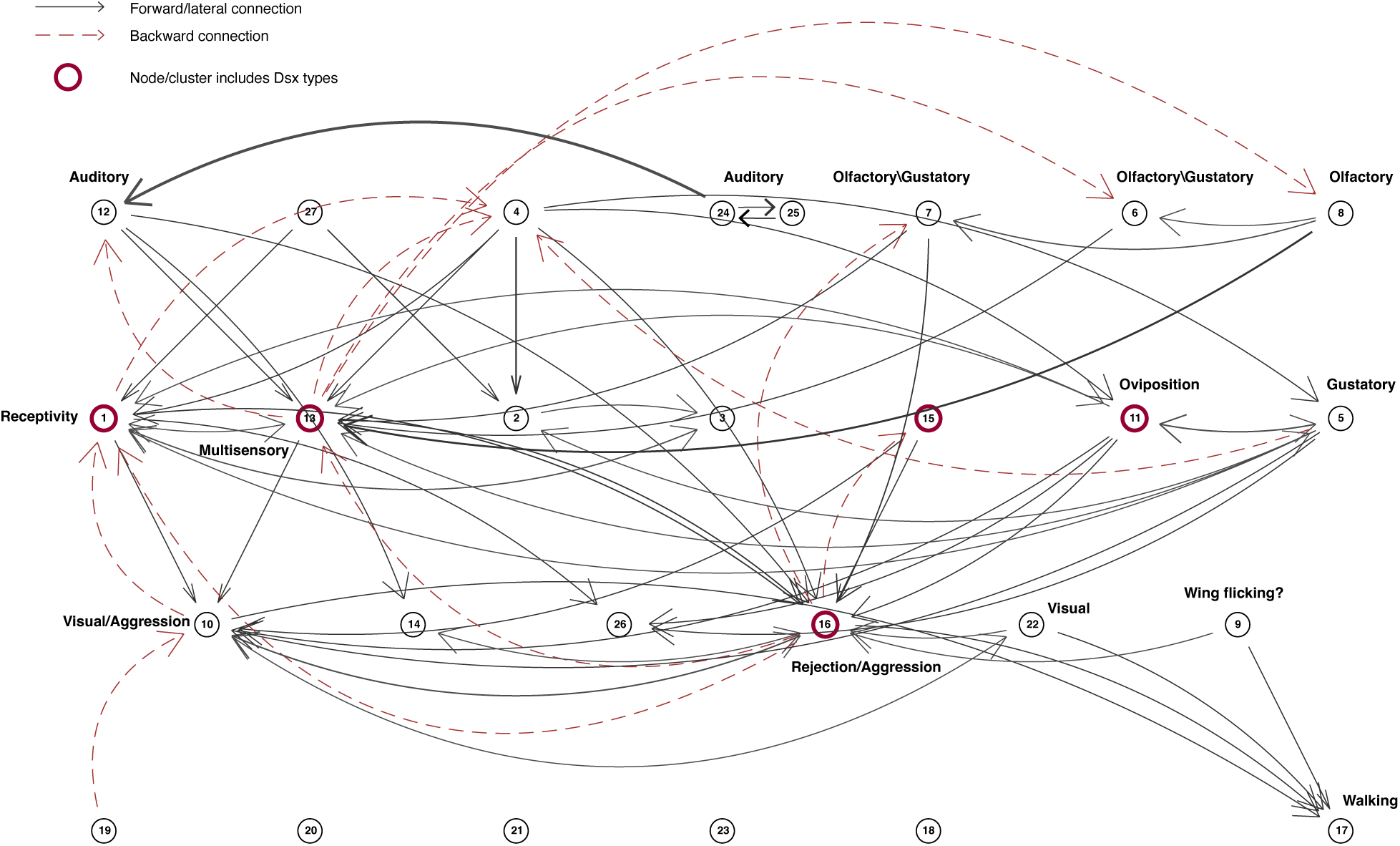
Network of significant connections between Fru/Dsx subtypes. Network diagram for Fru/Dsx clusters. Each Fru/Dsx cluster (Fig. 5) is represented as a single node, while edges represent normalized connectivity between clusters. Edge weights are calculated as the product of the fraction of output from node A that targets node B and the fraction of inputs to node B that originate from node A. Nodes were arranged into layers using a greedy algorithm to minimize backward arcs, followed by within-layer ordering to reduce edge crossings. While all weights were used to determine node positions, only the strongest 10% of edges are shown (see Methods). We infer the functional roles of some nodes based on the relative positions of their constituent cell types with respect to sensory modalities, as well as prior knowledge of the functions of specific cell types within each node. For example, cluster 8 (“olfactory”) includes two cell types, aDT3 and aDT11, both uniglomerular antennal lobe projection neurons. In contrast, cluster 16 (“rejection/aggression”) includes pMN1/DNp13, which controls ovipositor extrusion (a rejection behavior ^23,56^), and pC1d/e neurons, which regulate female aggression ^25,27^.

The most connected nodes are the clusters 1, 11, 13, and 16, all of which contain Dsx-expressing cell types. Together, connectivity-based clustering and network analysis (Figs. 5 and 6) provide an organizational layout of the Fru/Dsx neurons that should be useful for future functional work.

### Predicting Function from Connectivity

To make functional predictions, we focused on the connectivity around neurons driving opposing behaviors: vpoDN neurons, which respond to male pulse song and drive vaginal plate opening, *i.e*. receptivity ^21^, and DNp13 neurons, which respond to male pulse song ^17^ and drive ovipositor extrusion, *i.e.* rejection ^23,56^ (Fig. 7). We built the network in two steps starting from the strongest direct connections between these reception/rejection neurons and any of the Fru/Dsx neurons, then adding a second layer with a higher threshold (see Methods). We find that most of the Dsx cells are included in this network. Several recent studies had identified functions for these cell types using a mix of neural recordings, activation, or silencing experiments. pC1 neurons promote arousal ^21,25,27^, pC2l neurons are tuned to male courtship pulse song ^17,23,56^, and the SAG neurons drive female receptivity ^87^. Cell types selective for pulse song (pC2l, pC1d/e, vpoDN, DNp13 and aSP10) ^17,67^ are distributed into different subcircuits (Fig. 7A), suggesting how one mode of the male’s song can drive different behaviors, but it remains to be determined how the brain chooses between these different possibilities as courtship unfolds. There are four different subtypes of Dsx pC2l, and a Fru type, pIP5, with very similar morphology (Fig. 7B). The wiring diagram suggests that at least 3 pC2l types (pC2la-c) should respond to pulse song since they are postsynaptic to cell types tuned to pulse song (vpoEN ^21^ and aSP10 ^67^, Fig. 7C), while pIP5 should not. Using split-Gal4 lines (Supp Fig. 7 - S1) we recorded calcium responses to courtship song stimuli in these cell types (Fig. 7D-E, Supp Fig. 7 - S1; see Methods); consistent with our predictions, pC2la-d, but not pIP5 respond to male courtship song (Fig. 7D).

**Figure 7.**
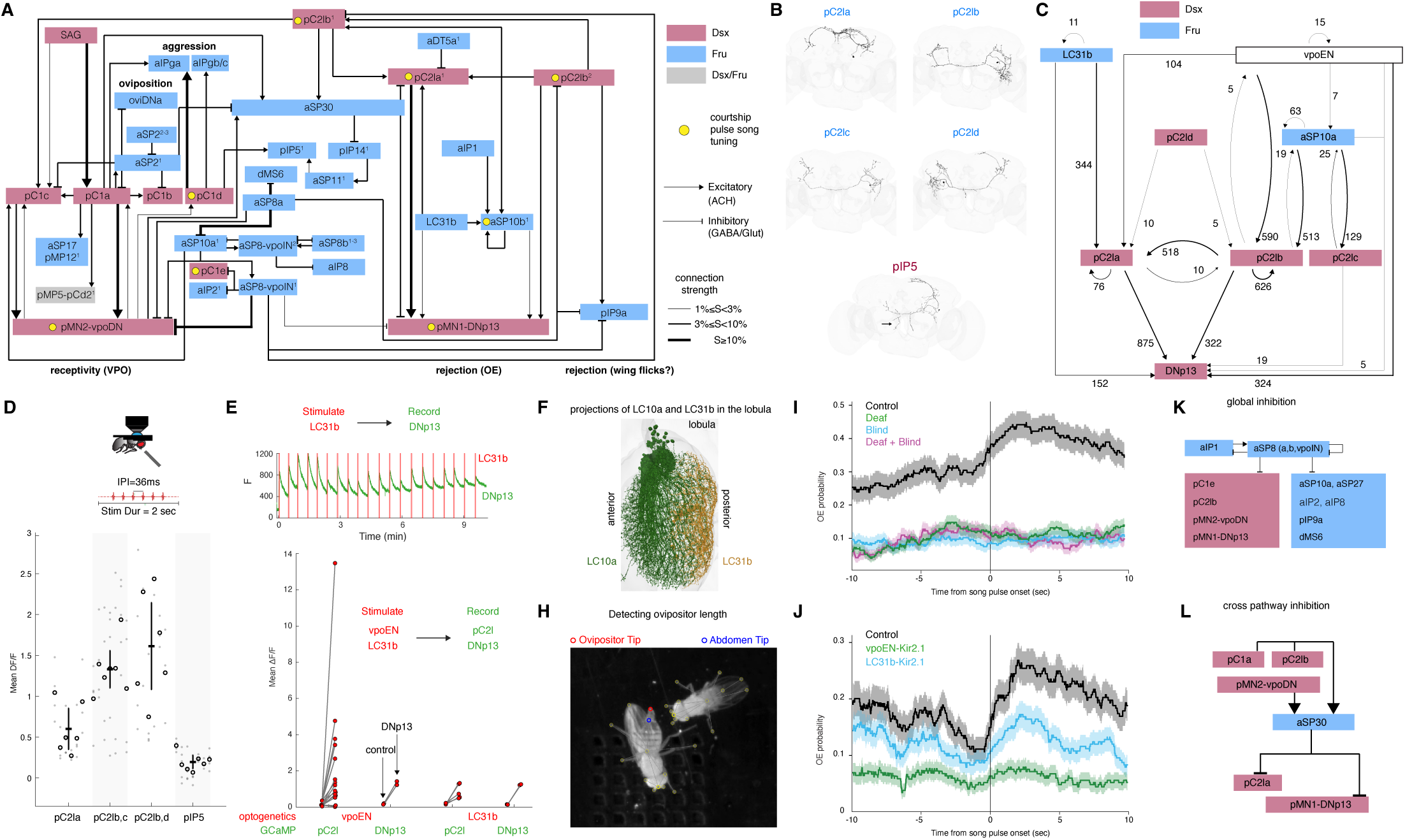
Predicting Function from Connectivity. **(A)** Strongest direct and indirect connections of pMN1-DNp13 and PMN2-vpoDN types with Fru/Dsx types. First, the subgroup of Fru/Dsx types with a strong (1% symmetric threshold; see Methods) directly connected to pMN1,2 were added. Then, the Fru/Dsx types connected to this subgroup, or cell types not in our Fru/Dsx list that are connected directly with pMN1-DNp13 or PMN2-vpoDN (using a 3% symmetric threshold) were added. Line width reflects connection strength (S), using three categories for the strength. Types that are known to be tuned to ‘Pulse song’ (one of the two major courtship song types in *D. melanogaster*) are marked with a red dot. None of the types in this diagram is known to be tuned to ‘Sine song’. Excitatory and inhibitory connections are represented by pointed and flat arrows, respectively. Connections are considered inhibitory if predicted in FlyWire as GABAergic or glutamatergic, and as excitatory if predicted as cholinergic, with the following caveat: SAG neurons are predicted to be Serotonergic in FlyWire but are shown to be cholinergic ^30^. ‘Fru’, ‘Dsx’ and ‘Fru or Dsx’ are colored in blue, red and gray as done elsewhere. Other types (e.g., AVLP340) are colored white. The asterisk near vpoEN is since this type is connected to other types in the diagrams and the arrows are eliminated for simplicity. Those connections are shown in Fig. 7C. **(B)** Dsx+pC2la-d and Fru+pIP5 have similar, but not identical morphologies. Black arrow points to lateral extension that exists in pIP5 but not in pC2la-d types. **(C)** A subset of the diagram in (A), showing interconnected pC2la-d types and adding in a non-Fru/Dsx pulse-tune auditory neuron vpoEN ^21^. This predicts that pC2la–d are all pulse-song responsive and that pC2l and DNp13 neurons integrate auditory input and visual input, via LC31b visual projection neurons (VPNs). While pIP5 has similar morphology as pC2la-d, it is not connected neither to vpoEN nor to pC2l types, therefore possibly not an auditory type. **(D)** To test whether the different pC2l and pIP5 types differentially respond to male courtship pulse song, we used split-GAL4 drivers that target each subtype. We found auditory responses in pC2la-d but not in pIP5. As pC2l neurons were already shown to be tuned to pulse song ^17,67^, while the auditory response of pIP5 neurons was not reported so far, for pC2l we tested only the response to pulse song, while for pIP5 we recorded the response to pulse song and to natural song bouts (Supp Fig. 7 - S1). Gray dot - single trials, circle - mean per fly. **(E)** We tested whether pC2l neurons and DNp13 neurons functionally respond to LC31b (visual) and vpoEN (auditory) activation (suggesting they are both multisensory) by expressing csChrimson in either LC31b or vpoEN using split-GAL4 drivers (see Methods and Supp Fig. 7 - S1), and imaging calcium responses in Dsx+ neurons. Responses were imaged in somas and cells were identified as either pC2l or DNp13 based on their spatial location. Control - a doublesex pC1 cell soma of the same fly and the same trial. Top: example trial: optogenetic stimulation of LC31b, Calcium response recorded in DNp13. **(F)** Innervation of the anterior and posterior lobula by example LC10a and LC31b cells - LC31b neurons likely detect visual motion behind the fly. **(H)** We tested whether OE (ovipositor extrusion behavior ^23,56^) was driven by multisensory interactions using SLEAP tracking (see Methods). **(I)** OE follows male pulse song onset in control females (black, n = 1788 OE events), but not in deaf (green, n = 1096), blind (blue, n = 1212), or deaf and blind (purple, n = 635) females. **(J)** OE following pulse song onset is reduced in either LC31b or vpoEN silenced females (control/vpoEN silenced/LC31b silenced, n = 2502, 1827, 2242 OE events, respectively). **(K)** aSP8 inhibits both pMN1-DNp13 and pMN2-vpoDN and Fru/Dsx cells that are strongly connected to those two types, therefore possibly driving ‘global inhibition’ in the rejection and receptivity. **(L)** aSP30 receives excitatory inputs from pMN2-vpoDN (which controls a receptivity female behavior, vaginal plate opening) and from other cells types which are strongly connected to pMN2-vpoDN, and sends inhibitory connections to pMN1-DNp13 (which controls a rejecting female behavior, ovipositor extrusion) and to cells that are strongly connected to pMN1-DNp13, therefore driving ‘cross inhibition’. aSP8-vpoIN^1^ = AVLP008, CB1385; aSP8-vpoIN^2^ = LHAV4c2; aSP8b^1–3^ = AVLP010, AVLP011,012; aIPgb/c = CB2131, CB1127; pC2la^1^ = CL313; pC2la^2^ = CB2375; pC2lb^1^ = AVLP567; pC2lb^2^ = AVLP569; pIP14^1^ = AOTU062; pSP7^1^ = SLPpm3_H01; pMP5-pCd2^1^ = SMP286;aDT5a^1^ = CL122_a; aSP10a^1^ = CB4244; aSP10b^1^ = CB1090, CB1783; aSP10b^2^ = CB3313; aIP2^1^ = CB1904, CB2593; pMP12^1^ = DNp60, pIP5^1^ = SIP201f, aSP11^1^ = AOTU008c, aSP2^1^ = SMP093, aSP2^2-3^ = SMP098_a, CB1456.

While prior work ^23,56^, suggested that ovipositor extrusion (OE), a rejection behavior, is driven by pulse song (via DNp13 neurons), we find from this analysis, that OE should be driven by *both* auditory and visual stimuli. To test this prediction, we optogenetically activated either vpoEN (auditory) or LC31b (visual) neurons and recorded from Dsx cell types implicated in rejection (pC2l and DNp13 ^56^), finding strong functional connections (Fig. 7E and Methods). LC31b neurons innervate only the posterior portion of the lobula (which receives information from the posterior portion of the visual field) making them prime candidates for processing male motion cues behind the female (Fig. 7F). This contrasts with LC10a neurons, implicated in female aggression ^86^, whose inputs are concentrated in the anterior lobula (aggression behaviors are driven by frontal motion cues). To further test if female flies combine spatially localized song and visual cues to drive rejection behaviors via this circuit, we measured OE during natural courtship behavior (males paired with mated females, see Methods, Fig. 7H). Either deafening the female or blinding her abolishes the OE response to courtship pulse song (Fig. 7I); thus, both cues are needed for OE. Silencing either vpoEN or LC31b neurons significantly reduces pulse-triggered OE (Fig. 7J), with silencing vpoEN having the larger effect (see Methods), suggesting there may be visual neurons other than LC31b also important for OE.

We also find additional motifs through analyzing the circuitry in Fig. 7A. vpoIN neurons (known to encode pulse songs outside of the conspecific range ^21^ including agonistic male song ^92^) provide global inhibition to a large number of Fru/Dsx neurons (Fig. 7K), not just vpoDN, as previously studied ^21,92^. This global inhibition ^93^ may sharpen tuning for conspecific courtship song in other auditory neurons but may also inhibit social and sexual behaviors when male agonistic song or male songs of another species are present ^94^.

Finally, we predict the function of aSP30 neurons, not previously characterized (Fig. 7L). 26% of the input synapses to aSP30 are from the neurons that putatively respond to the male pulse song (pC2lb-AVLP567), and 15.8% are direct inputs from the receptivity circuits (pC1a and vpoDN combined). aSP30 inhibits neurons in the rejection and aggression promoting circuits: 35% of its output synapses are with pC2la, pMN1-DNp13 and pC1d/e. This cross pathway inhibition motif forms the basis for a behavioral hierarchy ^95^ - when the receptivity pathway is activated, other behavioral circuits are shut down, giving receptivity a primacy. We do not uncover inhibition in the other direction (from rejection and aggression promoting circuits to the receptivity ones). Taken together, the diagram suggests two major pathways for the control of receptivity and aggression in *Drosophila* females, where multisensory signals, global and cross inhibition shape the chosen behavior.

## DISCUSSION

In this study, we provide a comprehensive mapping of putative *Fruitless* (Fru) and *Doublesex* (Dsx) neurons in the female *Drosophila* whole-brain connectome, identifying 98 Fru/Dsx types comprising over 1700 neurons. By combining light microscopy (LM) datasets with the FlyWire electron microscopy (EM) reconstructions, we establish a morphologically defined dataset of putative sexually dimorphic neurons and examine their connectivity patterns across the brain. This work offers insight into how Fru/Dsx circuits are embedded within broader neural networks and enables predictions about their function based on connectivity.

### Limitations of the Current Analysis

While extensive, our analysis captures only a subset of Fru/Dsx neurons in the female brain. Several neuron classes remain underrepresented or excluded, such as Kenyon cells of the mushroom body (*fru P1* is expressed in the mushroom body γ and αβ KCs ^14,96,97^), optic lobe neurons with ambiguous Fru expression ^39,60^, and sensory neurons whose morphology does not allow reliable Fru/Dsx classification. The neurons included in our dataset are chosen solely on the basis of their morphological overlap with LM images, and some neurons in our list may not express the *Fru* or *Dsx* genes. We divided the 98 Fru/Dsx types into subtypes, based on morphology and connectivity, and the fraction of input/output synapses each subtype has with any other Fru/Dsx neuron (Supp Fig. 1 - S3). Based on the observation that Fru/Dsx neurons tend to connect to each other, the fraction of connections with the Fru/Dsx network can serve as an indicator for which subtypes, within a given type, are more likely to be be Fru/Dsx expressing, or at least - to be part of the circuits involved in controlling social behaviors in flies.

### Comparisons with Analysis of the Male Whole-Brain Connectome

A recent study by Berg et al. (2025) performed a comprehensive comparison between the female connectome (FlyWire/FAFB, used here) and a new male brain connectome ^34^. Comparing both morphology and connectivity across the two EM datasets, each cell was assigned one of three categories: sexually dimorphic (matched between male and female, but with differences in morphology/connectivity), sex-specific (unmatched between male and female) or isomorphic (matched). Separately, and similar to our approach, Berg et al. also annotated neurons as putative Fru- or Dsx-expressing based on comparisons to light microscopy data. They provide information for nearly all FlyWire/FAFB female brain neurons on its dimorphism (sexually dimorphic, sex-specific, or isomorphic) and putative gene expression (‘dsx’,’fru’,’coexpress’): they report 3308 Fru/Dsx neurons in the female brain out of which 523 (15.8%) are female-specific or sexually dimorphic. In the other direction, there are a total of 1149 neurons that are female-specific or sexually dimorphic, out of which 523 are listed as Fru/Dsx (45.5%).

The best comparison between our study and theirs, is to compare the Fru/Dsx lists directly. Of the 3308 Fru/Dsx neurons identified by Berg S. et al., 2089 cells are not included in our Fru/Dsx list - some of these are in categories we did not identify (e.g, . olfactory sensory neurons), but some we found no LM matches for (e.g., 243 central complex neurons). Of our list of 1702 neurons, 72% of the neurons overlap with the neurons identified as Fru/Dsx/Coexpress (based on LM-EM comparison) in the Berg et al. dataset. Given that prior work suggests 1500-3000 neurons ^10,11,57^ or roughly 2% ^58^ of the neurons in the female brain are Fru while a few tens are Dsx ^19,46^, we believe we have missed Fru/Dsx neurons, while Berg et al. may have provided an overestimate. Berg et al. primarily followed Cachero et al. (2010)^10^, whereas we follow Yu et al. (2010)^11^ and Liu (2012) ^12^ - but many cell types can be mapped between schemes. In several cases, this mapping is one-to-one (e.g., aSP-f in Cachero et al. corresponds to aSP5 in Yu/Liu). In other cases, a single type in one scheme corresponds to multiple types in the other. For example, aDT-e in Berg et al. includes cells we annotate as aDT3 and aPN1, and aIP-e includes aIP3a, aIP3b, and aSP9. The remaining 28% of cells in our list (not identified as Fru/Dsx in Berg et al.) consist mainly of Fru types described in Yu et al. (2010) and Liu (2012), including aDT9, aIP8, aSP3, aSP8c, aSP17, aSP18, pIP13, dMS6, and pSP5. Several of these were identified as sexually dimorphic by Berg et al. (e.g., aIP8, aSP8c, and aSP17). In addition, we include several Fru cell types with projections in the SEZ ^45^ that are not annotated in Berg et al. Those types were identified in Sterne et al. 2021 ^45^. Our study adopted a conservative approach that prioritizes minimizing false positives, which likely contributes to the differences between studies. This conservative strategy is reflected in the enrichment of neurons in our dataset that are defined by Berg at al. as sexually dimorphic or female-specific: while 15.8% of the neurons annotated as Fru/Dsx in Berg et al. were also annotated as sexually dimorphic or female-specific, this proportion increases to 30.02% in our dataset.

### Implications for Understanding Female Behavior

Sexually dimorphic behaviors in *Drosophila*, including courtship, receptivity, aggression, and egg-laying, rely on distinct but overlapping neural circuits ^25,92,98,99^. Prior work has focused more on male-specific behaviors, such as courtship chasing ^73,100–102^ and singing ^16,18,103,104^. By anchoring Fru/Dsx neurons within a complete female connectome, our work illuminates the wiring logic of female social behaviors and opens new avenues for targeted functional experiments. Ultimately, this work lays the foundation for uncovering the complete circuit mechanisms underlying sexual and social behaviors in *Drosophila*. It enables researchers to move beyond isolated circuit fragments and toward a holistic understanding of how sensory inputs are integrated by Fru/Dsx neurons, transformed through local processing, and relayed to descending motor pathways. As larger datasets, including multiple brains and sex-specific connectomes, become available, the hypotheses and network motifs described here will serve as a critical scaffold for understanding how evolution sculpts the nervous system to generate sex-specific behaviors.

## Methods

### Identifying Fru/Dsx cells in FlyWire

Matches for Fru/Dsx neurons were identified in FlyWire (version 783) based on morphological similarity to light microscopy (LM) images. LM sources included image stacks from sparse genetic driver lines and stochastic labeling using Fru-Gal4 or Dsx-LexA lines. These sources included images of single neurons from Yu et al. (2010) ^11^, Liu (2012) ^12^, Cachero et al. (2010) ^10^, FlyCircuit (2011) ^39^, and Nojima et al. (2021) ^38^. We also used LM images of previously characterized neurons from the literature (see Table 1) ^21,30,40–42,45,4610–12,21,30,38–41,45–47^. We used a 3D image stack (rather than a 2D Z-projection) when available: 3D image stacks for Cachero et al. ^10^ and FlyCircuit ^39^ were downloaded from Virtual Fly Brain (https://www.virtualflybrain.org/) and a subset of the image stacks from Yu at al. were obtained through personal communication with Prof. Jai Yu (University of Chicago). We performed alignment of the FlyCircuit images from the FCWB to the JRC2018U template space using navis-flybrains (https://github.com/navis-org/navis-flybrains/). The Cachero et al. images were already available in the JRC2018U template space. We generated Color Maximum Intensity Projections (ColorMIP) images from binarized versions of images from FlyCircuit and from Cachero et al. ^105^ in a new Python implementation (https://github.com/htem/FANC_auto_recon/blob/main/fanc/render_neurons.py). Then, we ran image-based matchings against the collection of ColorMIP images for FlyWire v783 (as in ^105^) to infer a ranked list of FlyWire candidate neurons for each LM sourse. We then manually reviewed the candidate matchings with automatically generated Neuroglancer links showing the FlyWire EM candidates together with the corresponding mesh extracted from the binarized version of the single neuron clones in Cachero et. al. ^10^ or skeletonized single cells from FlyCircuit (Chiang et al. ^39^) to arrive at the final set of matches for further analyses. Where relevant, we applied an additional comparison to the Yu et al. 2010 images. Because these LM reconstruction data have more background noise, they were used mostly as a secondary source of validation to refine our list. We used 3D mesh and tracing data from this collection by transforming it into the FlyWire coordinate space with the Natverse toolset ^106^ and plotted candidate neurons alongside the corresponding meshes/tracings. After visual inspection, putative Fru+ neurons from the FlyWire dataset that exhibited weak spatial overlap with their corresponding reconstructions from the Yu 2010 dataset were discarded from our final list. Lastly, some of the electron-microscopy matches were first identified in the hemibrain ^48^, and then mapped to FlyWire using FlyWire Gateway (https://flywiregateway.pniapps.org/hemibrain_neurons), which was also used for finding candidates in the contralateral hemisphere. The Gateway candidates were manually sorted by comparison to the light microscopy images.

Using all sources described above, once FlyWire matches were identified, we used NBLAST ^43^ to identify more candidates. To further extend our search we used similarity in connectivity (using cell type annotation from ^36^), aiming to find missing individual cells in cell types that were recognized as Fru/Dsx.

Last, left-right symmetry was used both for perfecting the proofreading by identifying missing or extra branches on one hemisphere compared to the other one.

Nomenclature was largely adapted from Yu et al. (2010) ^11^ and Liu (2013) ^12^. The division into subtypes was done based on existing clustering into cell types that are largely defined by connectivity (^36,49^; see Table 1). It is likely that some subtypes or individual cells are Fru/Dsx negative (see Supp Fig. 1 - S3). Also, for 12 of 98 cell types we have lower confidence in the LM-EM match. This includes cases where the cell type was small and hard to distinguish (e.g., pIP7, aIP3a; Supp Fig. 2 - S2B) or when the neurons in LM or EM have a good but not perfect match (e.g., pIP14; Supp Fig. 2 - S1). Examining connectivity with the rest of the Fru/Dsx network suggests that of those 12 types, 10 have strong connectivity to other Fru/Dsx types (10% or more input or output synapses with Fru/Dsx types: aIP1,aIP3a,3b,aIP5, pIP12,14, pSP3,5, aSP5 and DN3A), while 2 have weak connectivity (pIP7,aSP13).

### Matched networks

In order to compare the connectivity of Fru/Dsx cells against populations of cells with similar properties we created 100 samples of ‘matched networks’. Matched networks were assembled as follows: (1) The same number of cells per superclass as in the Fru/Dsx group (Fig. 1C) was randomly chosen from all the FlyWire cells, excluding the Kenyon cells (as Fru/Dsx Kenyon cells were also excluded). (2) The pairwise euclidean distances between all the possible pairs of randomly selected cells was computed. If the mean distance was similar to the mean pairwise distance within the Fru/Dsx network (±2%), this group was selected as one a ‘matched network’; if not it was rejected. Distances are measures between cell bodies. (3) This procedure was repeated until 100 matched networks were selected.

Note that while the mean pairwise distance in the Fru/Dsx network is 210µm and the mean distance over groups of FlyWire cells (no kenyon cells, same superclass distribution) is 208.1 (<1% difference), most of the random networks are not within 2% of the Fru/Dsx mean pairwise distance due to the variability in the mean distances across random networks (±11.1µm).

### Measuring the bias for number of synapses with Fru/Dsx partners

Connections between proofread neurons are thresholded with 5 synapses for all the analyses described in this paper as previously done (e.g., ^35,107^). We aimed to reveal if Fru/Dsx cells are biased to have more synapses with a Fru/Dsx synaptic partner compared to the number of synapses they have with a random synaptic partner (Supp Fig. 1 - S5). We measured this bias separately for Fru/Dsx inputs and outputs. The procedure described here is for the inputs, a similar procedure was used for the outputs.

For each Fru/Dsx cell we made two calculations. First, we calculated the total number of input synapses it has with all its Fru/Dsx input partners (N_DsxFru, a single value). Second, we made 100 random choices of N input partners out of all the input partners of this cell (N being the number of Fru/Dsx input partners), and calculated the number of input synapses for this group (V_Rand, a vector with 100 values). Then we calculated the normalized bias (Supp Fig. 1 - S5) as (N_DsxFru-mean(V_Rand))/std(V_Rand), where ‘std’ stands for standard deviation. Positive/negative values mean a bias towards more/less synapses with Fru/Dsx partners, respectively.

### Rank analysis - information from sensory modalities

The traverse distance is a probabilistic measure for the number of hops from a group of neurons (the ‘seed’ neurons) to any individual neuron as defined in ^80^. The distance from the seed group to any other cell is calculated multiple times and then averaged. The average value is defined as the cells ‘rank’ with respect to the seed group. The distance is calculated probabilistically in the following way (as in ^80^). We start with an initial pool of cells called the ‘seed’ neurons. Then on each step, cells are added to the pool, where the probability P of a not-yet-traversed neuron to be added to the pool depends on the fraction of the inputs it receives from neurons already in the pool. For ‘fraction inputs’ between 0 and 0.3, P is the fraction divided by 0.3. For ‘fraction inputs’ of 0.3 or more, P=1. The number of steps it takes for a cell to be added to the pool is its traverse distance. This procedure is repeated multiple times (10000 times when we calculated the distance from sensory modalities and 1000 times when Fru/Dsx cell types were used as seeds) and then averaged the distance over repetitions. The ‘rank’ is calculated as 1+mean traverse distance, such that the cells in the seed group get a rank of 1 and the rest of the cells get a rank > 1. The cells included in each rank distance from 3 groups of sensory neurons (Mechanosensory-JO, gustatory, olfactory; ^35^) and from visual projection neurons ^35^ are shown in Fig. 4B . We normalized the rank to percentile to get a ‘normalized rank’ as in ^35^: the relative distance of a cell from the seed. For example, if a cell has a normalized rank of 20%, it means that 80% of the FlyWire cells have a larger rank from this Fru/Dsx type/seed (Fig. 4C-I). Small normalized rank means - shorter rank distance from the ‘seed’.

### Direct connectivity to descending neurons

First, connections between proofread neurons are thresholded with 5 synapses as for all the analyses described in this paper. Second, we identified all the descending types in FlyWire ^42^ that have at least 10% of their input synapses with Fru/Dsx neurons. For those descending neurons, any connection between a Fru/Dsx subtype and a descending type is shown. We used CODEX to find analogue cells between FAFB (v783) and BANC (v626). We show the BANC-predicted ^84^ effectors only when the effector is consistent across all individual cell types of that type. We only list the main body part that is predicted as an effector. For example, for bluebell., the effector Influence is “front_leg_neurosecretory_cell” and we denoted it in Fig. 8 as ‘leg’.

### Clustering of the Fru/Dsx subtypes

We used hierarchical clustering to group Fru/Dsx subtypes into connectivity clusters (Fig. 5). First, we calculated for each Fru/Dsx subtype *t,* a feature vector of length 2T, where T is the number of cell types in the FlyWire connectome. The first T values in the vector represent the number of synapses that cells of type t receive from every upstream type. The last T values in the feature vector count the number of synapses sent to each downstream type. We calculated distances between all pairs of Fru/Dsx subtypes using weighted Jaccard similarity. We chose weighted Jaccard similarity over alternatives like cosine distance and inverse L2 distance because those metrics either discard magnitude information altogether or are disproportionately dominated by the largest values (respectively), which is detrimental for accurately quantifying topological equivalence in sparse graphs. Experiments in ^49,108^ confirmed the superior accuracy of the Jaccard-based similarity measures for e.g. tighter clustering of same type neurons in the FlyWire conennctome. We applied hierarchical clustering with average linkage on the calculated distance matrix, and cut at distance 0.3 to define the connectivity clusters (see illustration in Supp Fig. 5 - S1).

### Connectivity diagram of the Fru/Dsx clusters

Connectivity diagram is shown between the 27 Fru/Dsx clusters (Fig. 6), where each cluster is represented as a node. Weights between pairs of nodes A, B is calculated as (fraction of all output synapses from A that are to B) X (fraction of input synapses to B that are from A). This way the weight is symmetric with respect to the input and output nodes. Sfas (small feedback arc set), an efficient implementation of a greedy algorithm for computing small feedback arc sets in directed weighted graphs (https://github.com/arie-matsliah/sfas?tab=readme-ov-file; ^109^) was implemented to position the 27 nodes in 4 layers. Then, node position within each layer was chosen to minimize edge crossing. While all weights are used for positioning the nodes, only the strongest connections (10% highest percentile) are shown.

## Experimental procedures - flies

Driver lines for vpoEN (SS61286) and pC2l (SS59882^21^, SS59848, SS59853) were made by Barry Dickson and Kaiyu Wang ^110^, all having a dsx-GAL4.DBD hemidriver. A driver line for LC31b (VT008646-p65.AD; R75A04-GAL4.DBD) was generated in the Murthy lab. A driver line for pIP5 (R91F05-p65.AD; fru.DBD) was generated in the Deutsch lab. Driver line used as LM resource in this study (see Table 1): pMP12: SS53468 (VT054914-p65.AD;VT040574-GAL4.DBD).

## Experimental procedures - microscopy

Adult female flies expressing GFP in Dsx pC2l, Fru pIP5 or LC31b neurons (Supp Fig. 7 - S1) were immunostained and scanned in a confocal microscope as described in ^17^. Fly preparation for two photon imaging, sound delivery, two photon imaging (Fig. 7D-E; Supp Fig. 7 - S1) and optogenetic stimulation were done as previously described in ^17,25^. DF(t) = F(t) - F_0_ was calculated by subtracting the mean baseline activity F_0_ (40% percentile of F in the 40 second window prior to stimulus onset) from F = F(t), the fluorescence signal in the green PMT channel. Mean(DF/F) was calculated over a 15 second window starting at the stimulus onset.

## Experimental procedures - behavior

For behavioral experiments, a single 4-5 days old virgin female was paired with a single wild type (NM91) male. For pairs that copulated within ∼1 hour, the male was removed 5-10 minutes after copulation was ended and the female was then paired with a naive wild type NM91 virgin male 18-24 hours later in a custom made copulation assay. Our behavioral assay was tiled with 9 pressure microphones and has a top-view camera as described elsewhere ^111^. ‘Sleap’ was used for pose-estimation ^111^, and the spatial-temporal profile of the distance between abdomen tip and ovipository tip was used for ovipository extrusion (OE) detection. Blinding was done by painting the female eyes with a black paint ^88^ and deafening was done by cutting both female aristae. Song segmentation was done using https://github.com/murthylab/MurthyLab_FlySongSegmenter as previously done ^25,88^ and OE probability was aligned to the onset of pulse song for each experimental condition.

For inactivation experiments Kir2.1, a hyperpolarizing inward rectifying potassium channel was used ^112,113^.

## Acknowledgements

We thank members of the Murthy and Deutsch laboratories for their advice on the project and comments on the manuscript; the staff at the Princeton Neuroscience Institute for support; and Google and the Allen Institute for assistance. DD was supported by the Zuckerman STEM leadership program and by the Israeli Science foundation (ISF grant 324/24); GSXEJ. by Wellcome Trust Collaborative Awards 203261/Z/16/Z and 220343/Z/20/Z, the Neuronex2 award (MRC_MC_EX_MR/T046279/1) and the MRC (MC-U105188491); and HSS and MM by NIH BRAIN Initiative grants RF1 MH117815, RF1 MH129268 and U24 NS126935.

## Author Contributions

*Project supervision*: DD and MM

*LM-EM matching and identifying Fru/Dsx types*: DD, AM, KW, AMo, AB, JH, JG, SG, GS, BD, AC

*Proofreading, annotation, and community support in FlyWire*: AMo, KW, SD, AB, JH, JG, S-CY, AS, CM, PS, MC, KE, YY, GSXEJ, BD, HSS

Experiments: EN, SJ, MW

*Connectivity-based analyses*: DD, AM, SD

*Writing*: DD, MM, and AC

---

**Table 1. Fru/Dsx cells**

**Table 2. Fru/Dsx clusters**

**Supp Figure 1 - S1.**
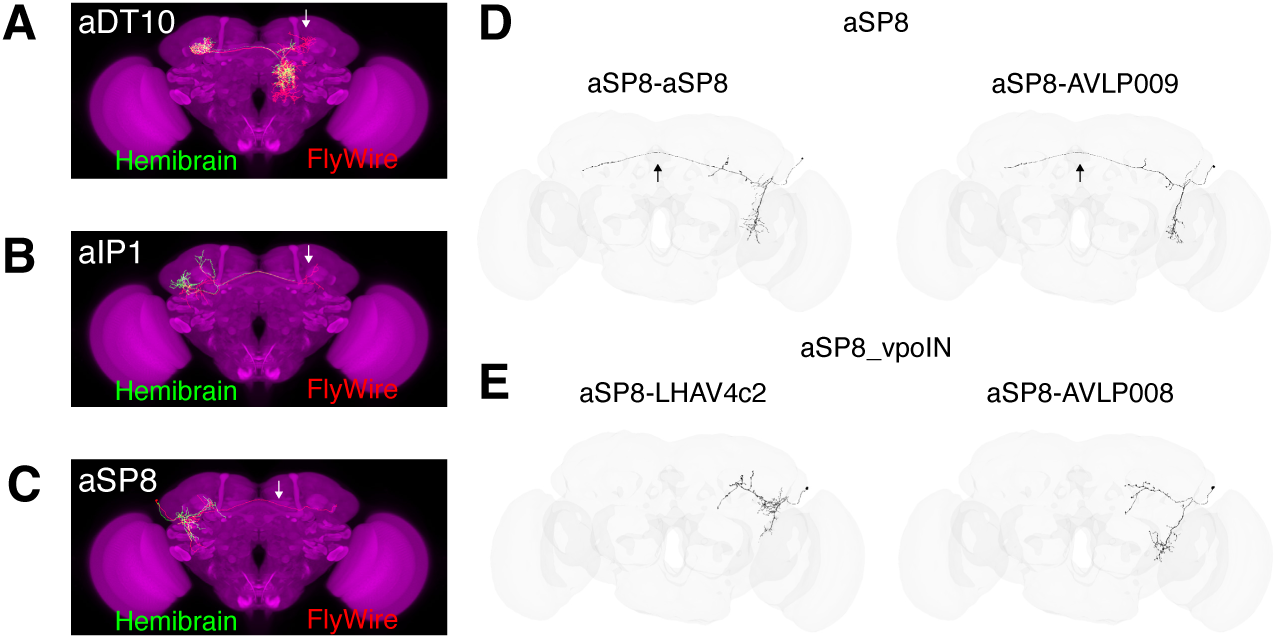
**(A)** Comparison of Fru/Dsx types in hemibrain and FlyWire wiring diagramsFru/Dsx aDT10 neurons in the FlyWire (^35^; red) and hemibrain (^48^; green) dataset. The two morphologies overlap, but the left lateral projection (ipsilateral to the soma; white arrow) is outside the hemibrain coordinates, and therefore missing in the aDT10 neurons in the hemibrain. **(B)** aIP1 in FlyWire and hemibrain datasets. The contralateral projection (white arrow) is missing in the hemibrain. **(C)** aSP8 in FlyWire and hemibrain datasets. The contralateral projection (white arrow) is missing in the hemibrain. **(D)** aSP8 and aSP8_vpoIN have overlapping morphologies, but aSP8_vpoIN neurons (previously annotated as vpoIN ^21^) are missing the contralateral projection (see black arrows). Based on this morphological difference, aSP8 and aSP_vpoIN are considered two distinct Fru/Dsx types. ‘aSP8’ and ‘aSP8_vpoIN’ have 2 subtypes each. Subtypes are based on a combination of morphology and connectivity ^36^ and are shown as cell types in CODEX ^44^.

**Supp Figure 1 - S2.**
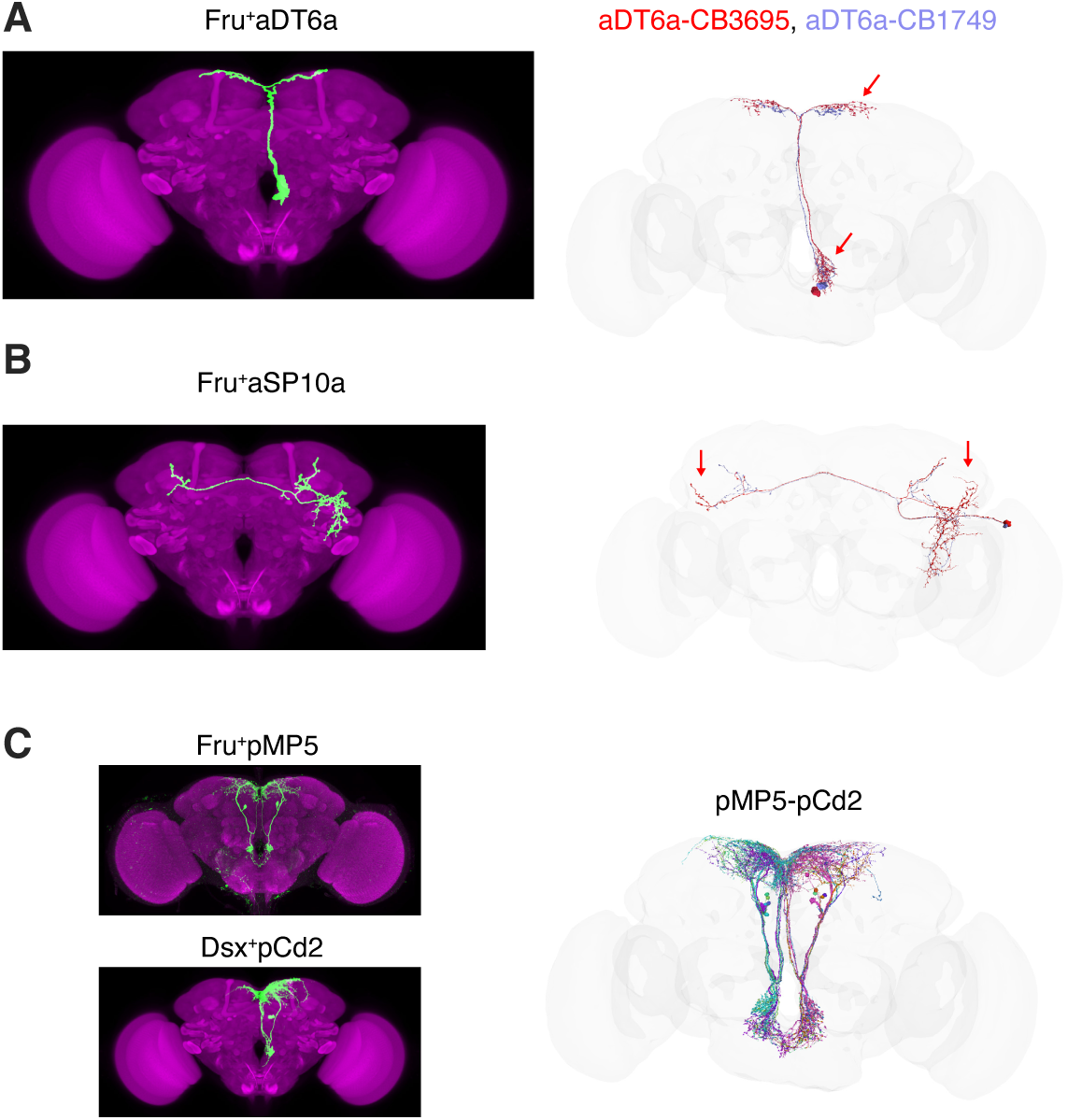
**(A,B)** Fru/Dsx types, defined by their morphology in light morphology images, can be composed of multiple subtypes in FlyWire. Subtypes may differ in subtle morphological details. **(A)** aDT6a, LM (left) and EM (right) images. Red arrows indicate dorsal-lateral projections and denser projections near the soma for aDT6a-CB3695 (red) compared to aDT6a-CB1749. aDT6a,b are heterogeneous populations that include both glutamatergic and cholinergic cells (see CODEX and ^7^). **(B)** extra projections in aSP10a-aVLP244 (blue arrows) compared to aSP10a-CB4244. Both subtypes match the LM image (left) well,**(C,D)** Fru+pMP3 has similar morphology as Dsx+pCd1, and Fru+pMN5 has similar morphology as Dsx+pCd2. Therefore, in this case, one group of FlyWire neurons potentially contains two Fru/Dsx cell types. **(C)** Similar morphology for pCd2 and pMP5. This type in FlyWire is termed pMP5-pCd2. Similarly, pC1d has similar morphology with pMP3, and both are termed pMP3-pCd1.

**Supp Figure 1 - S3.**
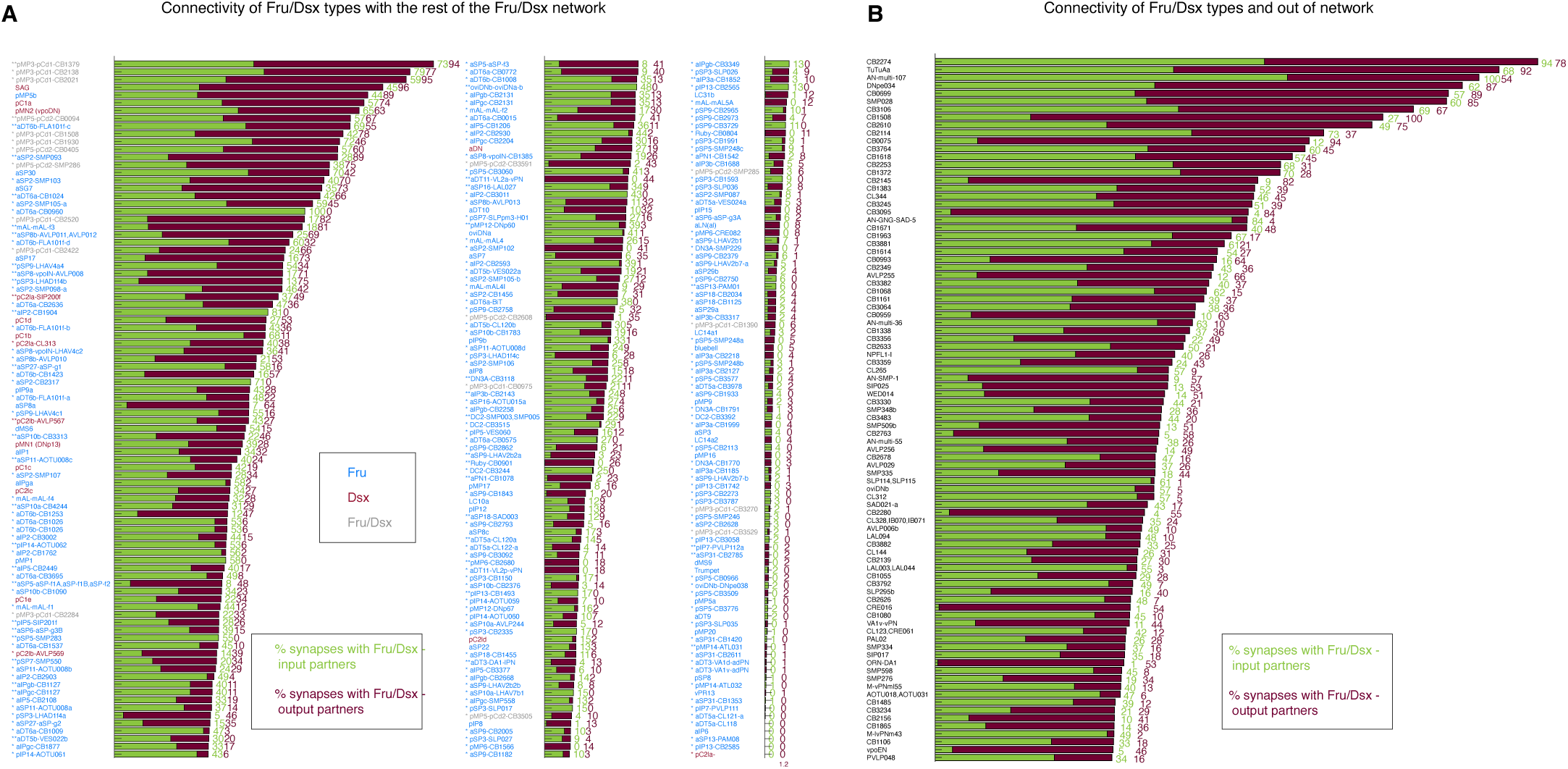
Connectivity in the Fru/Dsx network. **(A)** The percentage of synapses each Fru/Dsx subtype has with any Fru/Dsx partner (including within type), for inputs (synapses with presynaptic partners; green) and outputs (postsynaptic partners; magenta-red). When a cell type has multiple subtypes, * or ** appear near the subtype name. ** indicates that for a given type, this subtype has the most connections with other Fru/Dsx neurons. For example, pMP3/pCd1-CB3301 has 73% of input synapses from other Fru/Dsx neurons and 94% output synapses, more than the input+output % for any other subtype of pMP3/pCd1. When a given Fru/Dsx type has multiple subtypes but some cells do not have a defined primary type in Codex, the Fru/Dsx type is used also as the subtype name (e.g., ‘pSP9-pSP9’). This is in contrast to cases where there is only one primary type per Fru/Dsx type (e.g., pC1a or aSP30) that are not marked with anasterisk. The total number of synapses per type and the number of Fru/Dsx synapses per type both exclude within-subtype connections. **(B)** Same as (D), but for FlyWire cell types not in the Fru/Dsx list, that have a large proportion of connections with Fru/Dsx neurons (input+output = 50% or more). These cell types are candidates for being Fru or Dsx positive. The most-connected types in this list are the CB2274 neurons (morphologically similar to the pMP3-pCd1) and TuTuAa neurons (Supp Fig. 1 - S6,). TuTuAa was recently shown to be involved in enabling visually guided aggressive pursuit in females ^86^. CL062 neurons shown to drive male and female aggression ^114^ are also not labeled as Fru/Dsx. They make >10% of their inputs and outputs with Fru/Dsx cells including Dsx pC1d and Fru aIPg neurons, previously shown to drive female aggression ^25,27^ as well as with the Fru pMP12 (descending types DNp60, DNp67).

**Supp Figure 1 - S4.**
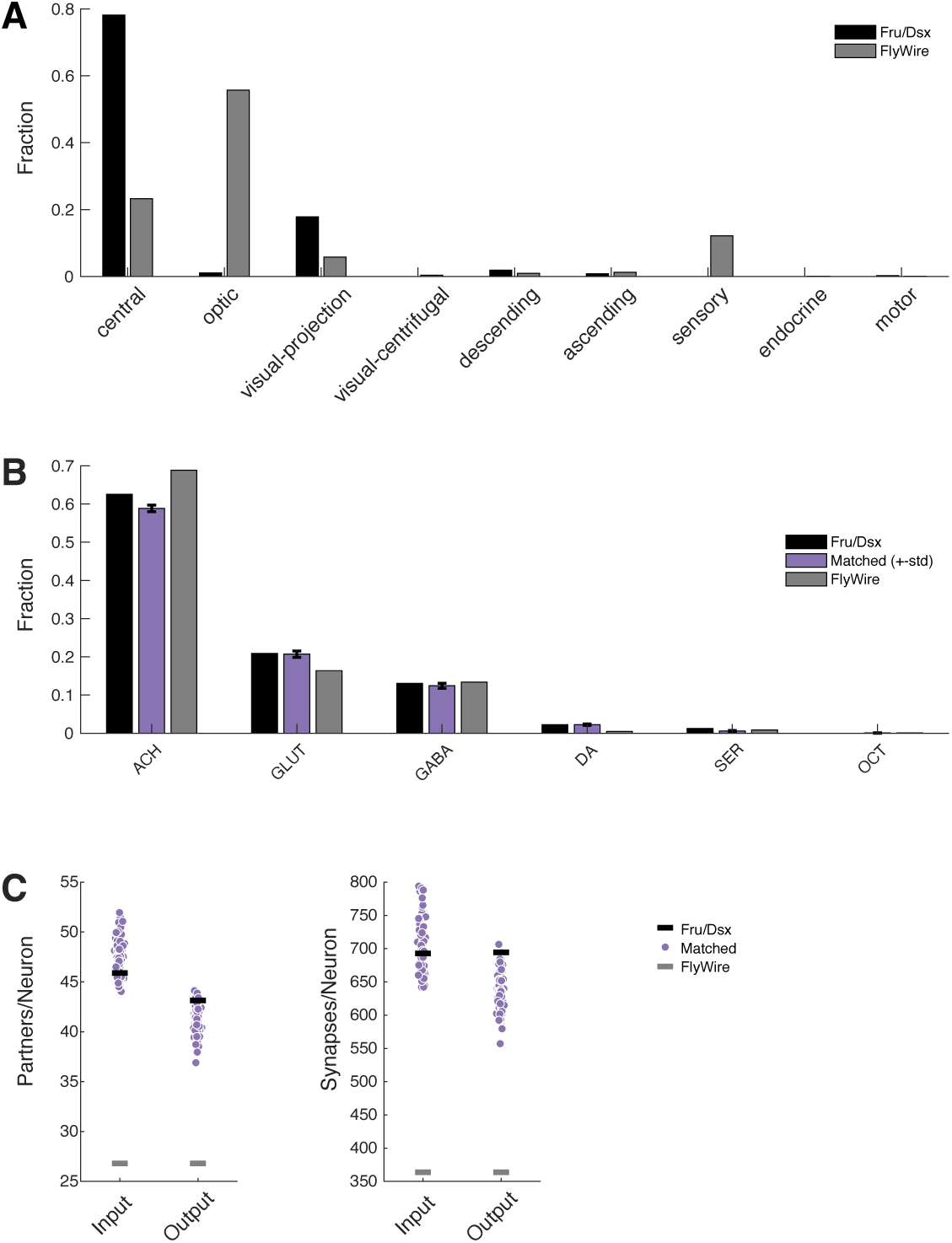
Comparing Fru/Dsx and ‘Matched’ Networks. **(A)** Distribution of Fru/Dsx neurons (black) and all FlyWire neurons (gray) across ‘superclasses’ ^35^. **(B)** Distribution of predicted neurotransmitters for all FlyWire neurons (gray), for Fru/Dsx neurons (black), and for neurons in the 100 matched networks (purple). **(C)** Mean number of synaptic partners per neuron (left) and number of synapses per neuron (right) for Fru/Dsx neurons (black), all FlyWire neurons (gray) and for each of 100 matched networks (purple).

**Supp Figure 1 - S5.**
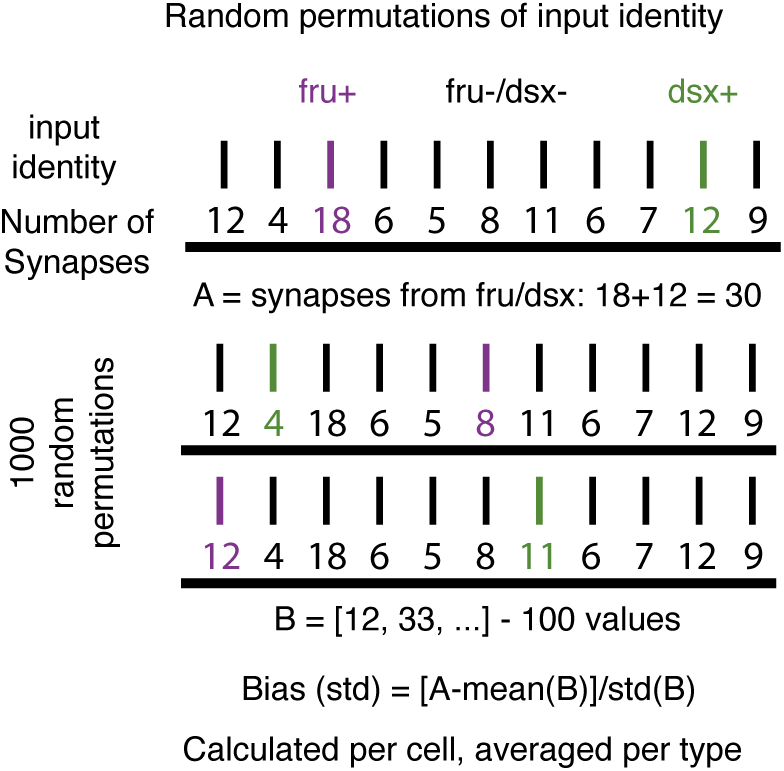
Calculating the bias for having more or less synapses with Fru/Dsx partners. Illustration of the calculation done in Fig. 1J (see Methods for details). The tendency of Fru/Dsx neurons to have more or less synapses with Fru/Dsx partners compared to the number of synapses with random partners is calculated separately for input and output synapses (here illustrated only for the inputs). In short, for each cell in the Fru/Dsx list, the total number of synapses with Fru/Dsx neurons is calculated (30 in this example). Then the identity of the partner neurons as being Fru+ (blue), Dsx+ (red) or not Fru/Dsx (black) is shuffled 1000 times, and each time the total number of synapses is calculated for the ‘Fru’/’Dsx’ partners. Then, the bias is calculated as the difference between the real number of the synapses with Fru/Dsx partners (A), minus the mean number of synapses with Fru/Dsx partners (B), divided by the standard deviation of B. The average bias is calculated over all the cells in any Fru/Dsx type. The distribution of the biases over the cell types is shown in Fig. 1J.

**Supp Figure 1 - S6.**
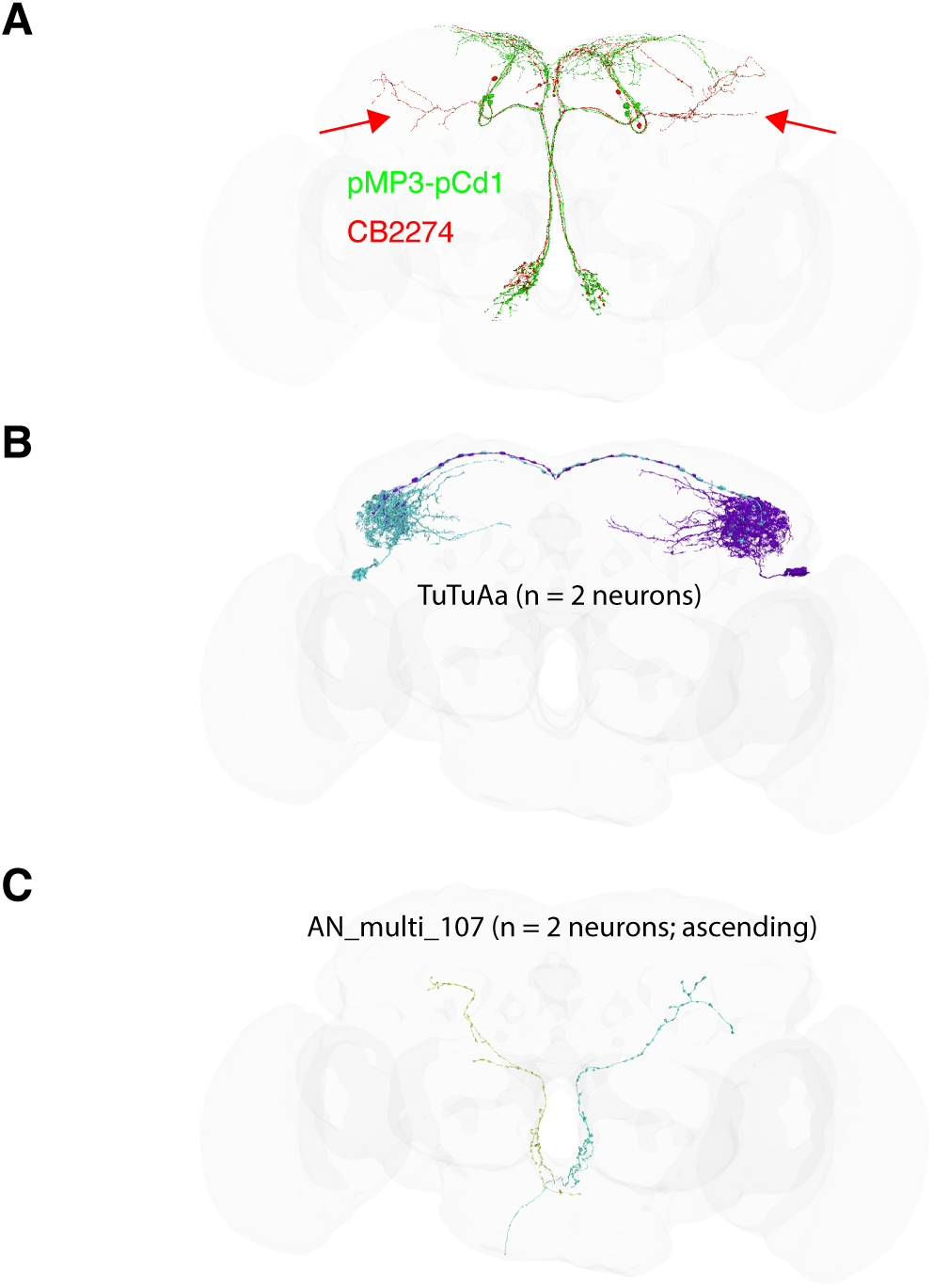
Cell types not listed as Fru/Dsx, with strong connectivity to Fru/Dsx cells. **(A)** cell type CB2274 has 94% and 78% of its input and output synapses with Fru/Dsx cells, respectively. These cells overlap in morphology with the Fru/Dsx type pMP3/pCd1. CB2274 has extra branches (red arrows) that are not present in pMP3/pCd1, therefore it was excluded from the Fru/Dsx candidate list. Given its strong connectivity with the Fru/Dsx network, those cells are possibly Fru/Dsx. **(B)** the glutamatergic TuTuA neurons that were previously shown to be involved in a circuit controlling female aggression ^86^ have 68% and 92% of their input and output synapses with Fru/Dsx cells. TuTuA neurons are downstream Fru aIPg and upstream Fru LC10 neurons, and are likely Fru expressing cells. **(C)** The ascending neurons type AN_multi_107 have 100% and 54% of their synapses in the brain with Fru/Dsx cells. As those are ascending neurons, their major input is at the VNC, an area missing from the FlyWire/FAFB dataset.

**Supp Figure 2 - S1.**
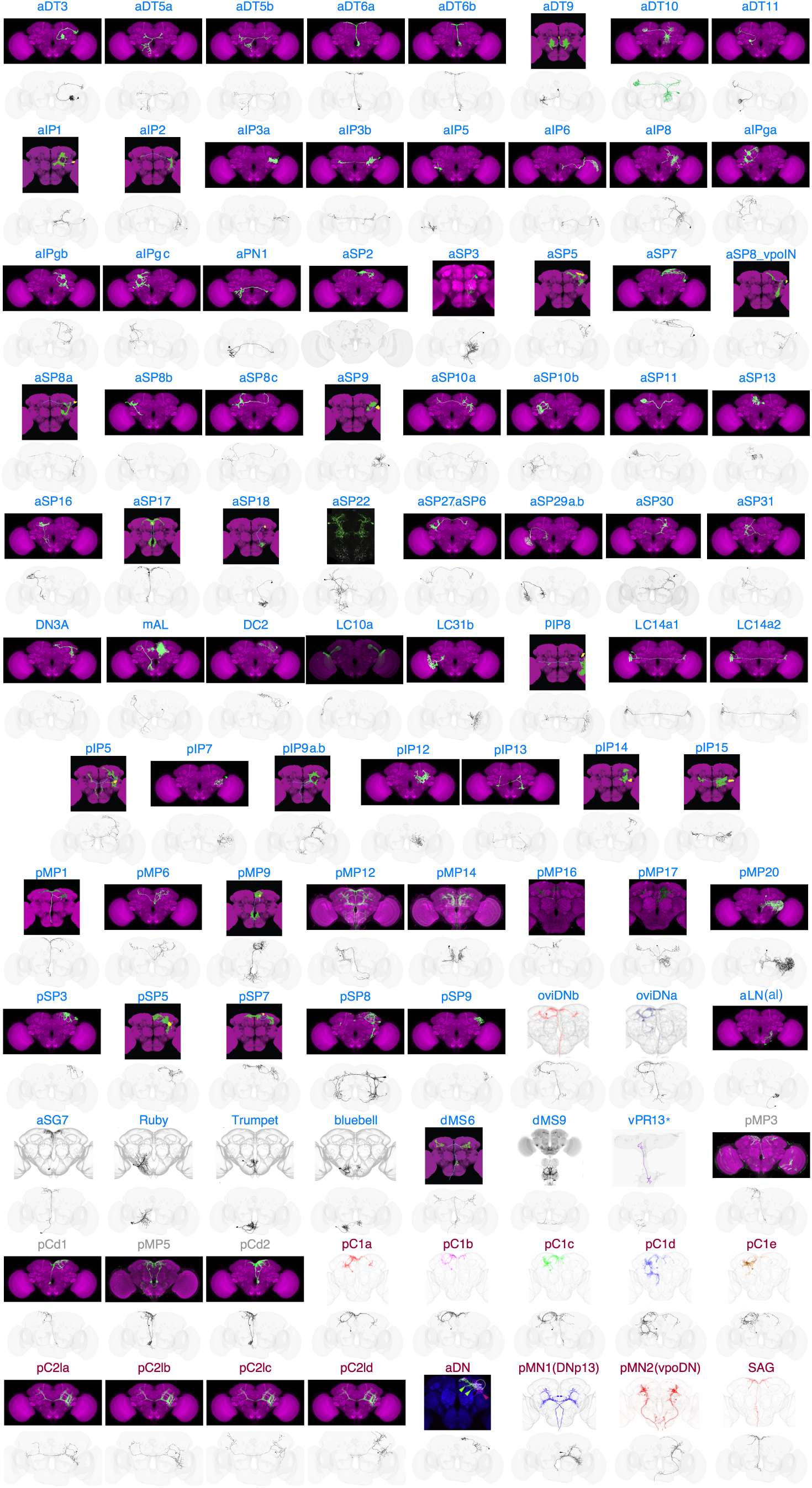
LM-to-EM matches for each Fru/Dsx type. For all the 98 Fru/Dsx types, a light microscopy (LM) image (top) and electron microscopy (EM) reconstruction from FlyWire (bottom) is shown. For EM images, we chose a single example cell. The LM source for each type and a FlyWire link corresponding to the displayed neuron are provided in Table 1. In a few cases a single LM image was used for multiple Fru/Dsx types: pC2l (pC2la, pC2lb, pC2lc, pC2ld) and LC14a (LC14a1, LC14a2), aSP27/aSP6, pIP9a (descending)/pIPb (central) and aSP29a (LC33)/aSP29b(LT51). Several Fru/Dsx types have additional subtypes (based on annotations in Codex) - see Table 1 and Supp Fig 1 - S3. Fru/Dsx types are ordered in alphabetical order (first Fru, then Fru/Dsx, and then Dsx). Cell type name is colored as elsewhere: Fru in blue, Dsx in red, and Fru-Dsx (Fru/Dsx types pMP3-pCd1 and pMP5-pCd2; for those types there is shared morphology between a Fru and a Dsx type) in gray. For cell type vPR13, the source image is not from LM, but rather a single cell from BANC (ID: 720575941404900464, Type: AN_GNG_SAD_32) that shares morphology with vPR13 and overlaps in the FlyWire/FAFB in the brain. This image is marked with *.

**Supp Figure 2 - S2.**
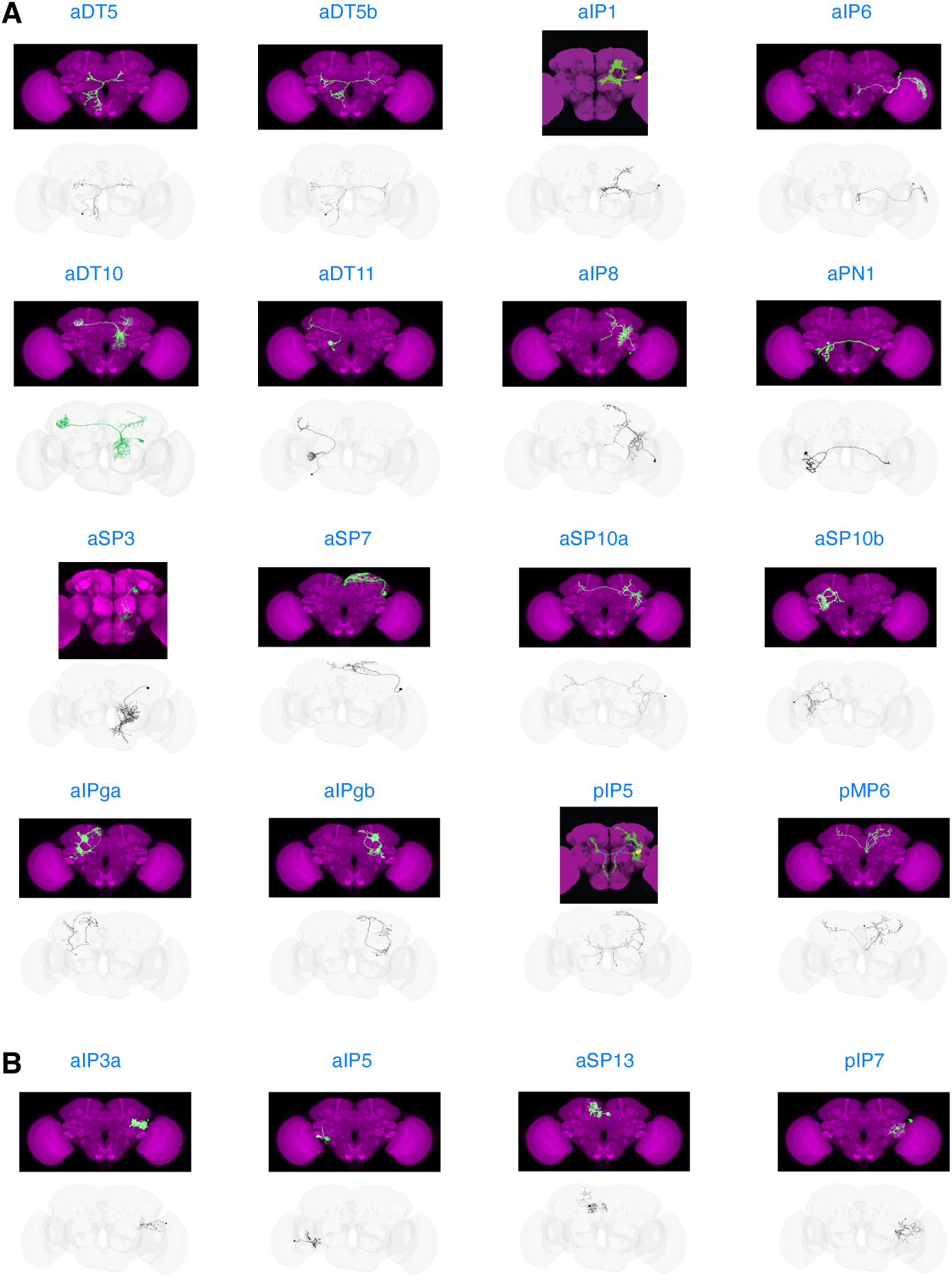
LM-to-EM matches for selected Fru/Dsx type. For 20 out of the 98 Fru/Dsx types, a light microscopy (LM) image (top) and electron microscopy (EM) reconstruction from FlyWire (bottom) is shown (all the 98 types are shown in Supp Fig. 2 - S1). For EM images, we chose a single example cell. The LM source for each type is provided in Table 1. **(A)** example cell types with clearly distinct morphology, for which we have high confidence about the LM-EM matches. **(B)** example cell types with less distinct morphology, for which we have a lower confidence in the LM-EM matches.

**Supp Figure 2 - S3.**
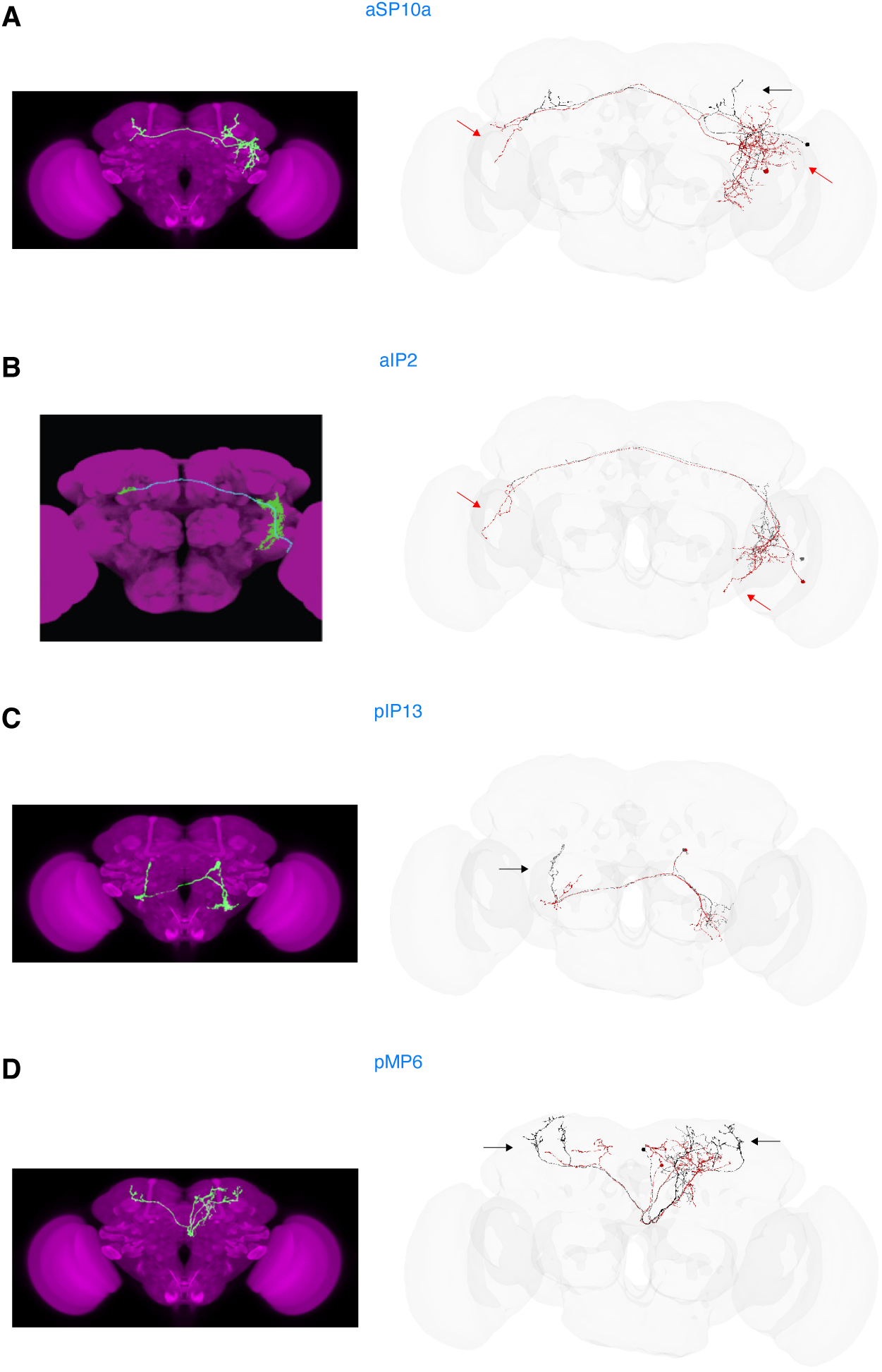
borderline cases: FlyWire cell with partial LM-EM overlap not considered as Fru/Dsx. Example of borderline cases in EM-LM matches. In those examples, cells with better overlap were considered Fru/Dsx (black), while cells with some mismatch with LM were not considered as Fru/Dsx (red). Black/red arrows indicate extra branches in the black/red example cells. **(A,B)** red aSP10a and aIP2 cells have extra branches that do not appear in LM, **(C,D)** red pIP13, pMP6 cells are missing branches that appear in LM.

**Supp Figure 3 - S1.**
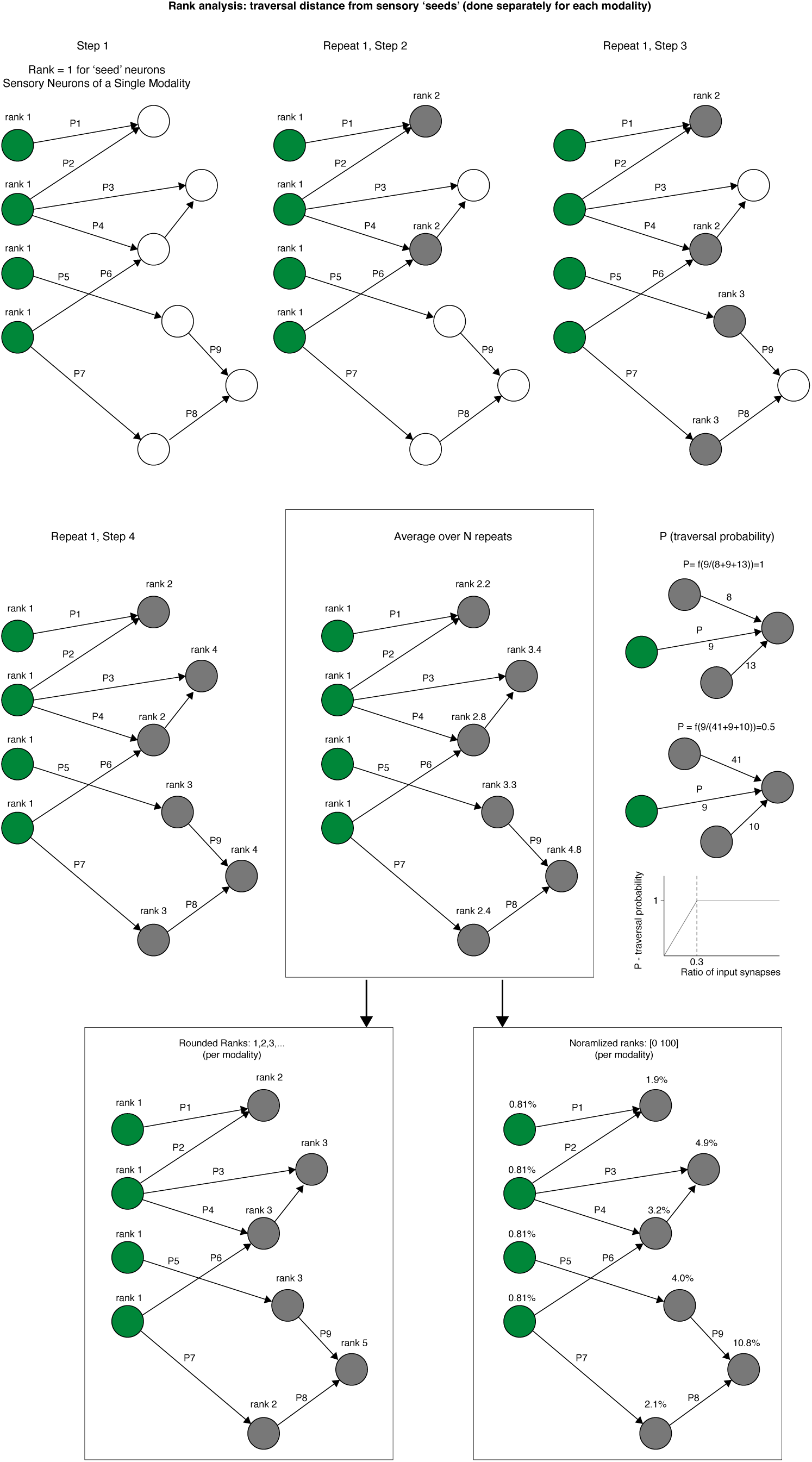
rank analysis – schematics. Transversal analysis is calculated as previously done ^35,80^. Starting from a seed (here, group of all sensory neurons of a given modality identified in flyWire/FAFB), a probabilistic model is used for propagation into other cells. At the first step, the ‘seed’ neurons get a ‘rank’ of 1. Then, in step 2, more cells are added, denoted ‘rank’ of 2 etc. This procedure is repeated N times. In each repeat, each cell in flyWire gets a rank. After N repeats the rank (or ‘transversal distance’) of each individual neuron is calculated as an average over N repeats. The rank is then either rounded (see Fig. 3B), or normalized by converting all the traversal distances to percentiles (therefore, in the range 0–100; See Fig. 3C-I). The normalized distances are also used for creating a 2D embedding, based on the rank distances of all the FlyWire cells from sensory modalities (see ^35^). The location of each Fru/Dsx neuron (excluding the optic and visual projection neurons; see Methods) is then projected onto this map (Fig. 3 and Supp Fig. 3 - S2)

**Supp Figure 3 - S2.**
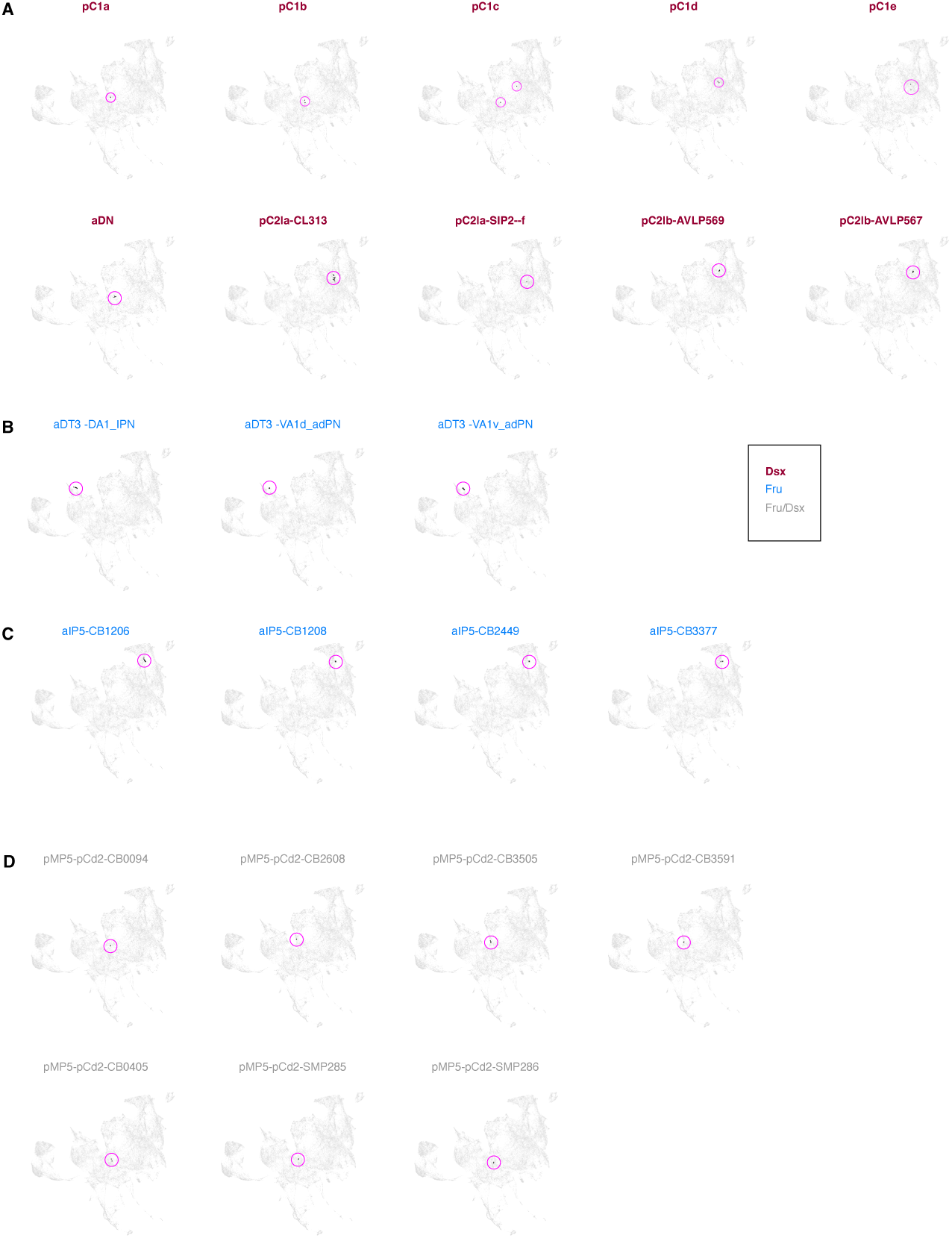
Embedding of Fru/Dsx neurons into UMAP embedding of the sensory space. The embedding is taken from Dorkenwald et al., 2024 ^35^. For each type,all individual cells are shown as black dots. The pink circles are for visualization purposes only. (A) Dsx types., (B) Fru types, (C) Fru-Dsx types. Colors for Dsx/Fru/Fru-Dsx are chosen as elsewhere.

**Supp Figure 3 - S3.**
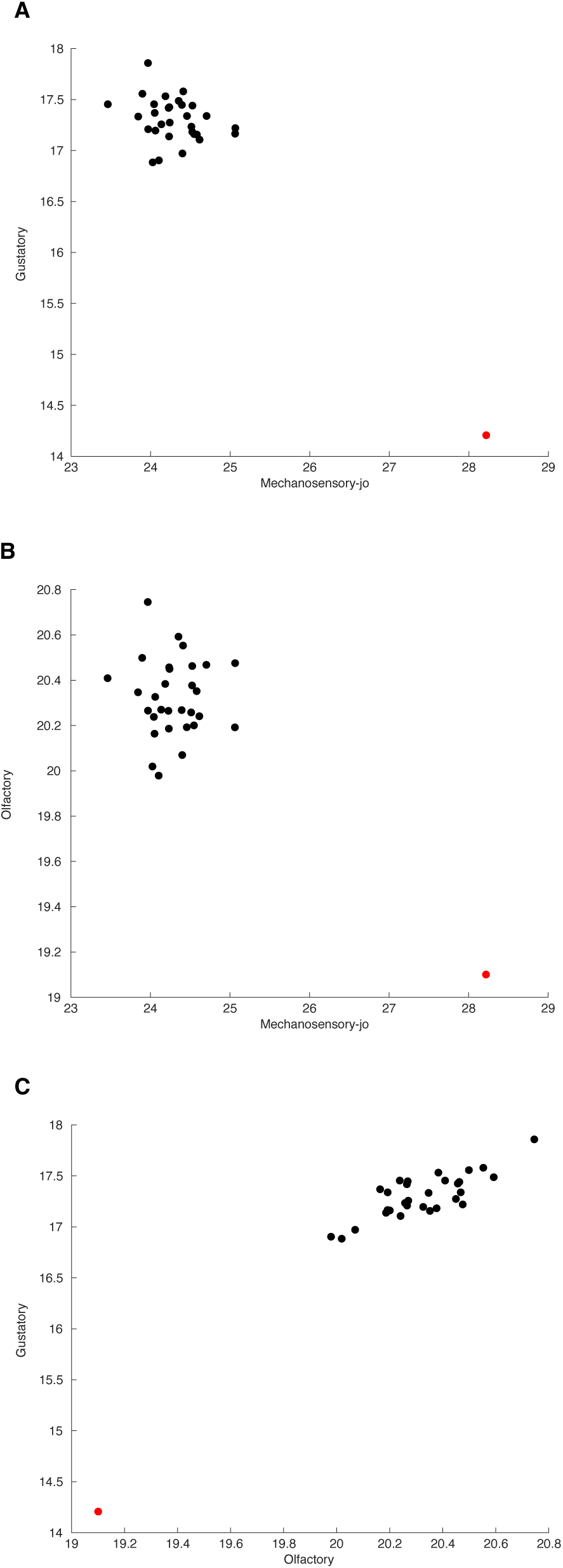
multisensory integration. Normalized traversal distances from pairs of modalities.

**Supp Figure 4 - S1.**
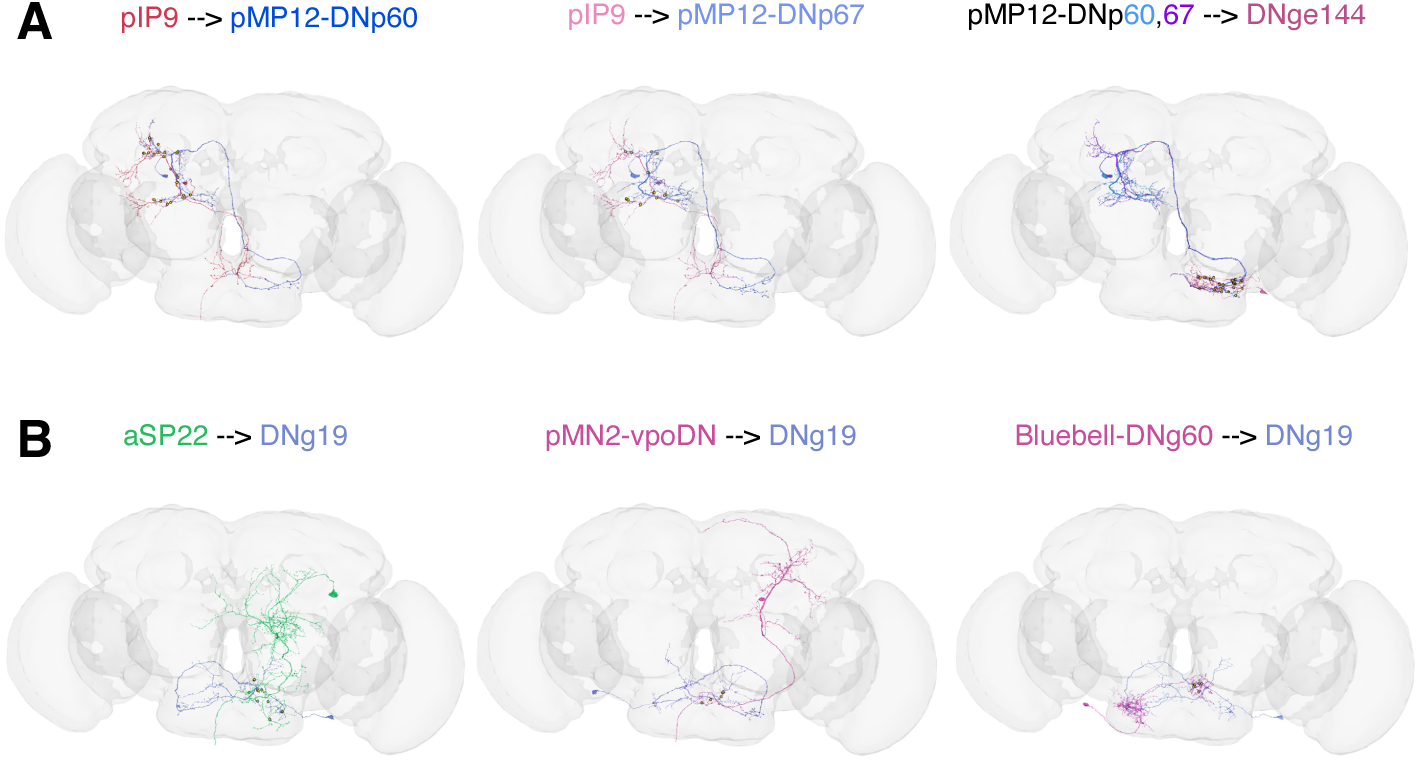
**(A)** Examples for connections between pairs of descending cells: fru+pIP9 is upstream of both pMN12-DNp60 (left), and pMP12-DNp67 (middle). Both pMP12-DNp60 and pMP12-DNp67 are directly upstream of DNge144, a pair of descending neurons (one per hemisphere) that are not in the Fru/Dsx list. Synapses are shown as red circles. **(B)** Three Descending types have strong direct connections upstream the descending cell type DNg19 (which is not in the Fru/Dsx list): fru+aSP22 (left), dsx+pMN2-vpoDN (middle) and fru+Bluebell-DNg60. Synapses are marked in yellow.

**Supp Figure 5 - S1.**
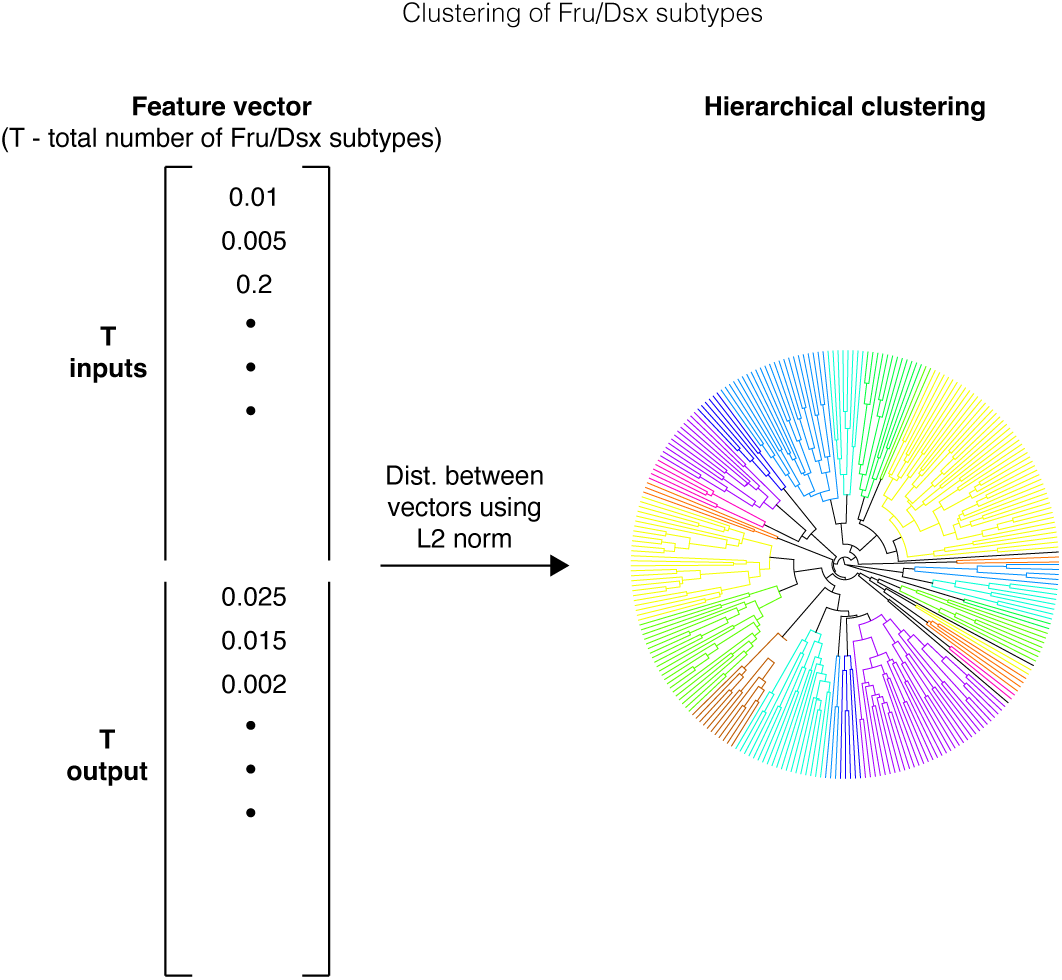
clustering by similarity in connectivity. we calculated for each Fru/Dsx subtype *t,* a feature vector of length 2T, where T is the number of cell types in the FlyWire connectome. The first T values in the vector represent the number of synapses that cells of type t receive from every upstream type. The last T values in the feature vector count the number of synapses sent to each downstream type. We calculated distances between all pairs of Fru/Dsx subtypes as L2 norm between their corresponding feature vectors and applied hierarchical clustering with average linkage on the calculated distance matrix, and cut at distance 0.3 to define the connectivity clusters (see Fig. 5).

**Supp Figure 6 - S1.**
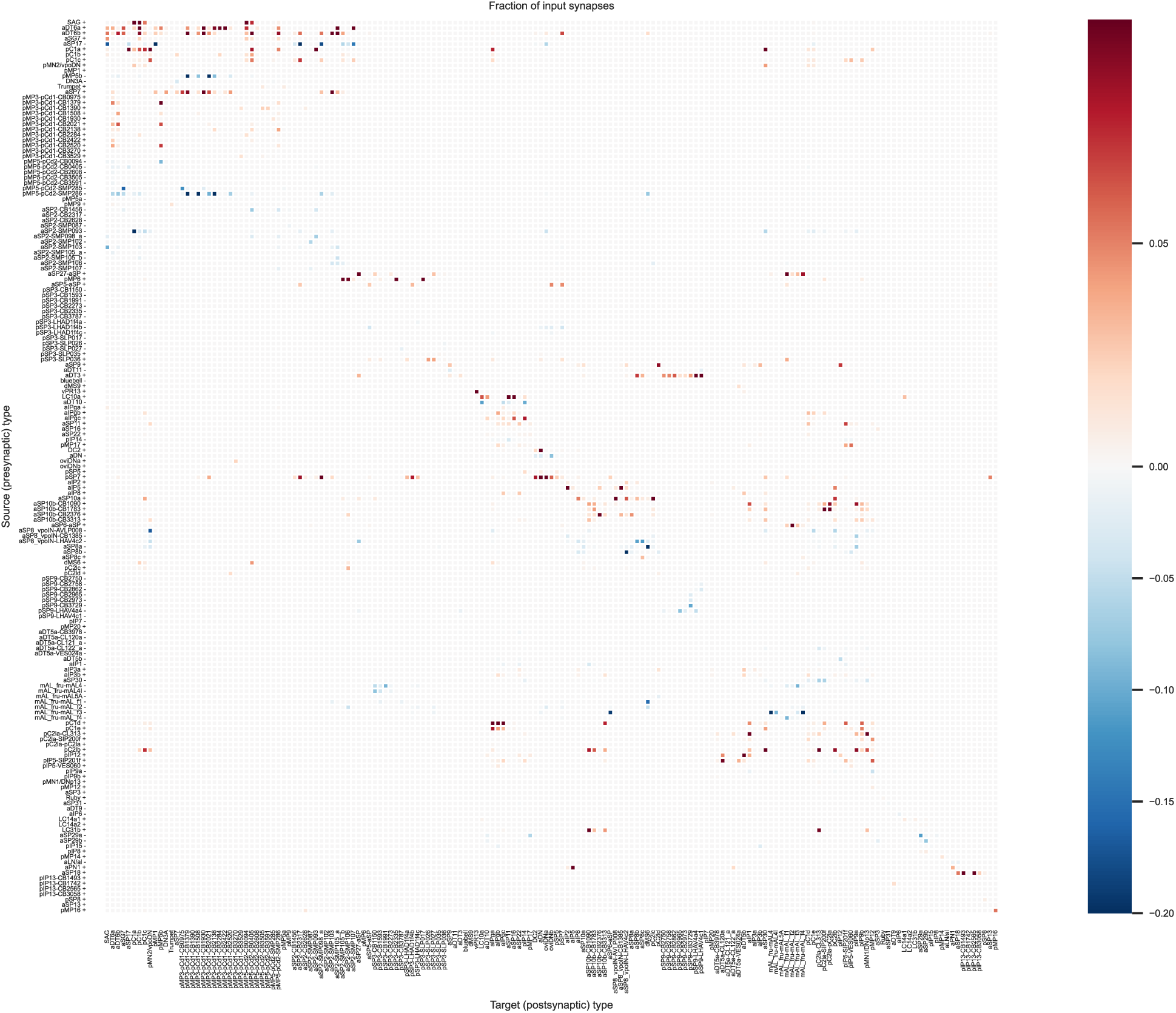
fraction of input synapses. For each Fru/Dsx type (target) the fraction of input synapses with each Fru/Dsx type (source) is shown (see Methods).

**Supp Figure 6 - S2 -.**
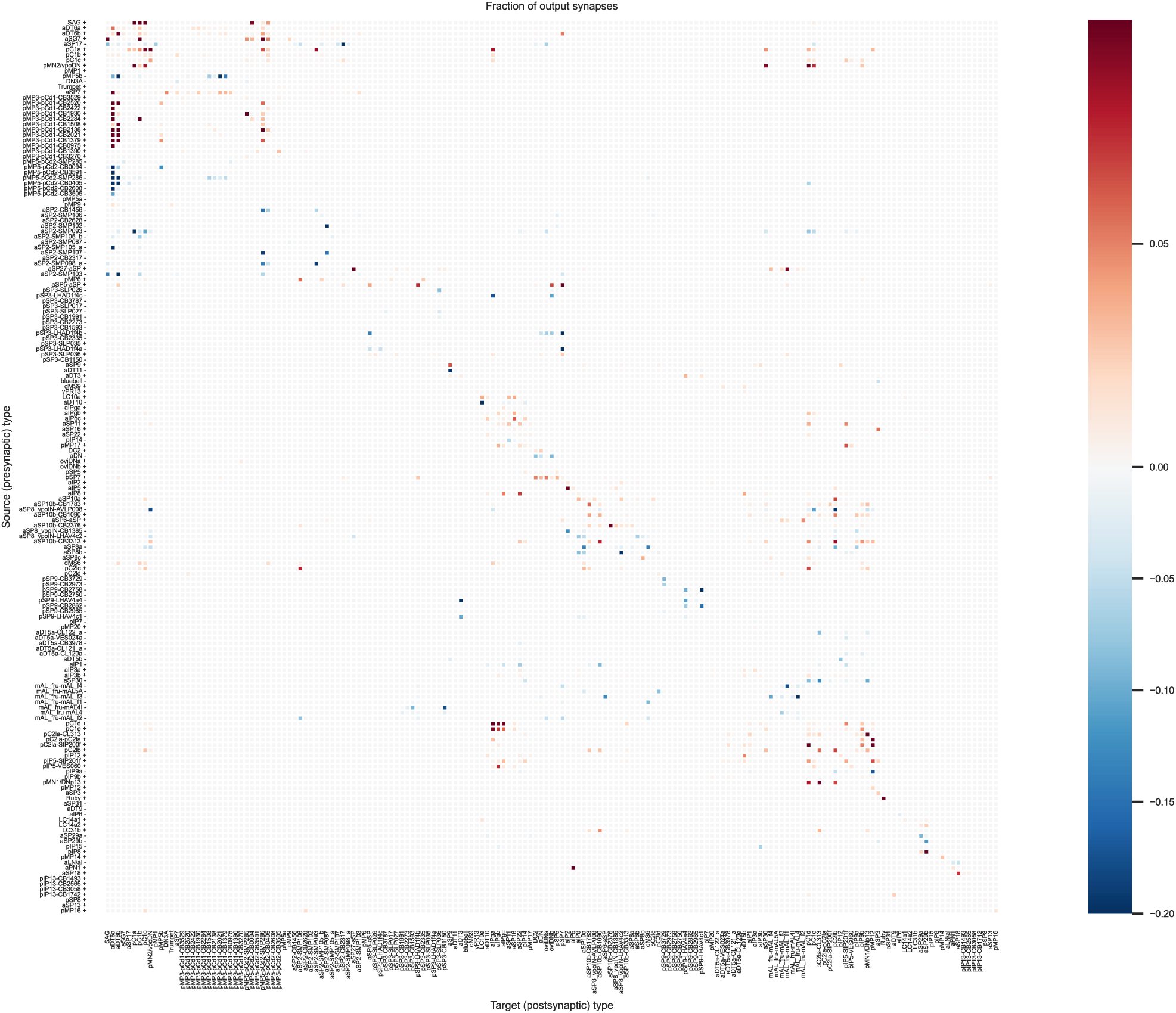
fraction of output synapses. For each Fru/Dsx type (source) the fraction of output synapses with each Fru/Dsx type (target) is shown (see Methods).

**Supp Figure 7 - S1.**
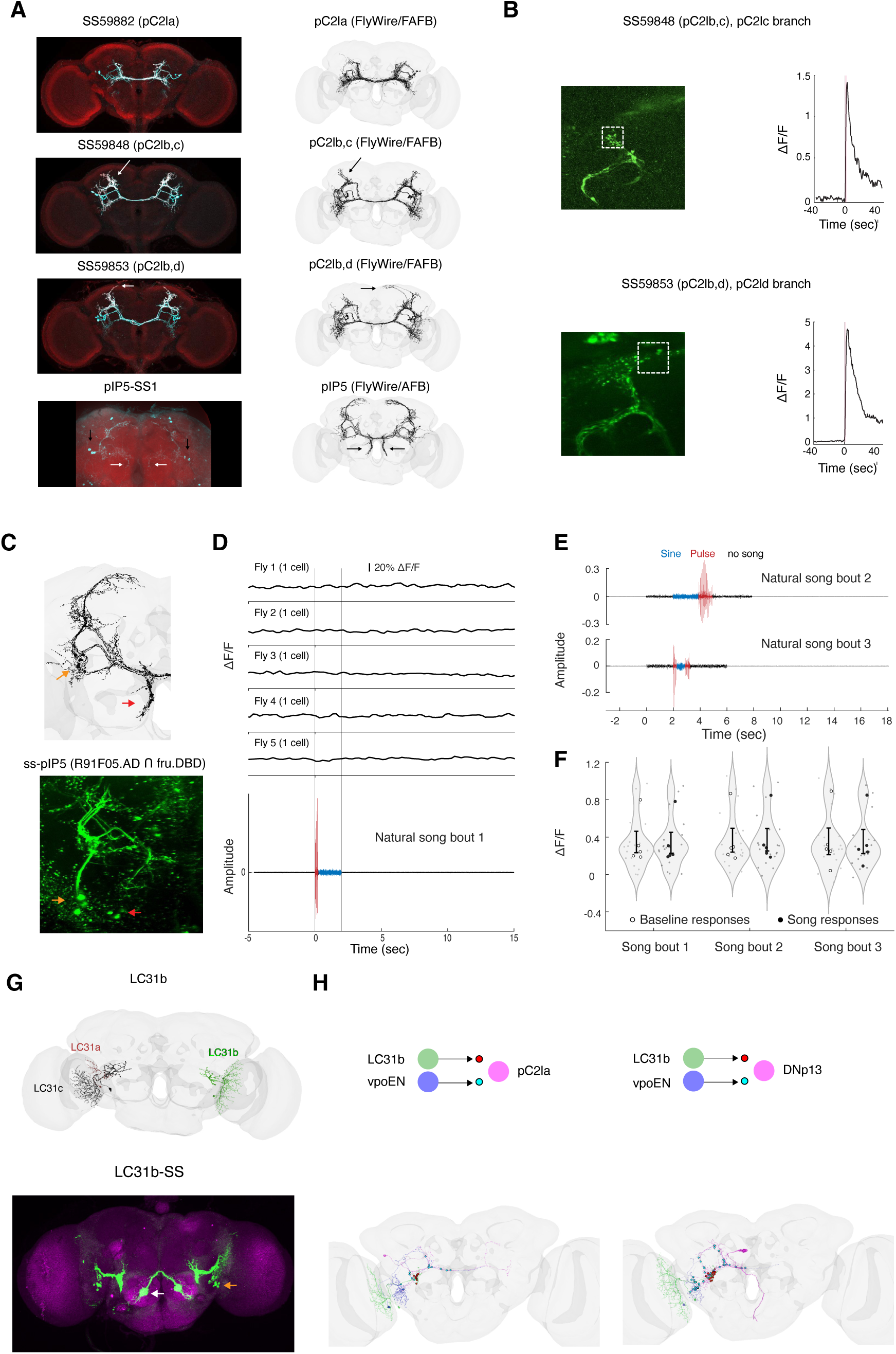
**(A)** GFP labeling of dsx+pC2l and fru+ pIP5 cells using split-gal4 driver lines (see Methods) and corresponding cells in FlyWire.. White arrow for SS59848 indicates arborization that exists in pC2lc, but not in pC2lb. White arrow for SS59853 points to arborization that appears in pC2ld but not in pC2lb. Corresponding black arrows point to the same neural process in FlyWire. In the z-projection of pIP5-SS1, the black arrows point to cell bodies (4.4±0.72 cells per hemisphere; n = 5 flies), while the white arrows point to projections near the midline that appear in Fru pIP5, but not in Dsx pC2l cells. **(B)** Calcium response in specific branches. Top: Driver line SS59848 likely contains pC2lb,c cells. Response shown for the area marked as white dotted box. This branch exists in pC2lc but not in pC2lc neurons. Bottom: SS59853 (likely containing pC2lb,d), response in a branch that exists in pC2ld, but not in pC2lb. **(C)** pIP5 in flyWire (top) and under a two-photon microscope (bottom). Orange arrows: cell bodies. Red arrows: medial branch that exists in pIP5 but not in pC2l. **(D)** Example Calcium responses of 5 single cells (1 per fly) for neutral song bout. Bottom: Natural song bout number 1 **(E)** Natural song bouts 2,3. **(F)** DF/F before (‘baseline’) and during stimulus presentation in response to 3 song bouts. Small gray dots represent single trials. Large dots represent per-fly averages (empty: baseline; full: during song bout). **(F)** Synapses to a single pC2l neuron (top) and to a single DNp13 neuron (bottom) from a visual projection neuron (LC31b; red circles represent synapses) and auditory neuron (vpoEN; cyan circles represent synapses). **(G)** Top: LC31a,b,c in FlyWire. Bottom: GFP labeling of LC31b. Orange arrow: cell bodies. White arrow: GFP signal that is not from LC31b neurons. **(H)** Schematics (Top) and FlyWire images for the convergence of the Pulse-responsive cell type vpoEN (synapses in Cyan) and the visual projection neurons LC31b (synapses in red) onto pC2la and DNp13.

## References

1. Rideout, E. J., Billeter, J.-C. & Goodwin, S. F. The sex-determination genes fruitless and doublesex specify a neural substrate required for courtship song. Curr. Biol. 17, 1473–1478 (2007).

2. Dickson, B. J. Wired for sex: the neurobiology of Drosophila mating decisions. Science 322, 904–909 (2008).

3. Kimura, K.-I., Hachiya, T., Koganezawa, M., Tazawa, T. & Yamamoto, D. Fruitless and doublesex coordinate to generate male-specific neurons that can initiate courtship. Neuron 59, 759–769 (2008).

4. Wohl, M., Ishii, K. & Asahina, K. Layered roles of fruitless isoforms in specification and function of male aggression-promoting neurons in Drosophila. Elife 9, (2020).

5. Rideout, E. J., Dornan, A. J., Neville, M. C., Eadie, S. & Goodwin, S. F. Control of sexual differentiation and behavior by the doublesex gene in Drosophila melanogaster. Nat. Neurosci. 13, 458–466 (2010).

6. Brovkina, M. V. & Clowney, E. J. Fruitless produces diverse neuronal sex differences by editing lineage-programmed chromatin landscapes. *bioRxiv* (2026) doi:10.64898/2026.04.14.718444.

7. Allen, A. M., Neville, M. C., Nojima, T., Alejevski, F. & Goodwin, S. F. Differential neuronal survival defines a novel axis of sexual dimorphism in the Drosophila brain. Cell Genom. 6, 101125 (2026).

8. Zhou, C., Pan, Y., Robinett, C. C., Meissner, G. W. & Baker, B. S. Central brain neurons expressing doublesex regulate female receptivity in Drosophila. Neuron 83, 149–163 (2014).

9. Kimura, K.-I., Ote, M., Tazawa, T. & Yamamoto, D. Fruitless specifies sexually dimorphic neural circuitry in the Drosophila brain. Nature 438, 229–233 (2005).

10. Cachero, S., Ostrovsky, A. D., Yu, J. Y., Dickson, B. J. & Jefferis, G. S. X. E. Sexual dimorphism in the fly brain. Curr. Biol. 20, 1589–1601 (2010).

11. Yu, J. Y., Kanai, M. I., Demir, E., Jefferis, G. S. X. E. & Dickson, B. J. Cellular Organization of the Neural Circuit that Drives Drosophila Courtship Behavior. Curr. Biol. 20, 1602–1614 (2010).

12. Liu, T. Anatomical Functional Dissection Fruitless Positive Courtship Circuit Drosophila Melanogaster. Na. (2012).

13. Brovkina, M. V., Duffié, R., Burtis, A. E. C. & Clowney, E. J. Fruitless decommissions regulatory elements to implement cell-type-specific neuronal masculinization. PLoS Genet. 17, e1009338 (2021).

14. Palmateer, C. M. et al. Single-cell transcriptome profiles of Drosophila fruitless-expressing neurons from both sexes. Elife 12, (2023).

15. Clowney, E. J., Iguchi, S., Bussell, J. J., Scheer, E. & Ruta, V. Multimodal Chemosensory Circuits Controlling Male Courtship in Drosophila. Neuron 87, 1036–1049 (2015).

16. von Philipsborn, A. C. et al. Neuronal control of Drosophila courtship song. Neuron 69, 509–522 (2011).

17. Deutsch, D., Clemens, J., Thiberge, S. Y., Guan, G. & Murthy, M. Shared Song Detector Neurons in Drosophila Male and Female Brains Drive Sex-Specific Behaviors. Curr. Biol. 29, 3200–3215.e5 (2019).

18. Roemschied, F. A. et al. Flexible circuit mechanisms for context-dependent song sequencing. Nature 622, 794–801 (2023).

19. Rezával, C. et al. Activation of Latent Courtship Circuitry in the Brain of Drosophila Females Induces Male-like Behaviors. Curr. Biol. 26, 2508–2515 (2016).

20. Rezával, C. et al. Neural circuitry underlying Drosophila female postmating behavioral responses. Curr. Biol. 22, 1155–1165 (2012).

21. Wang, K. et al. Neural circuit mechanisms of sexual receptivity in Drosophila females. Nature 589, 577–581 (2021).

22. Kerwin, P., Yuan, J. & von Philipsborn, A. C. Female copulation song is modulated by seminal fluid. Nat. Commun. 11, 1430 (2020).

23. Mezzera, C. et al. Ovipositor Extrusion Promotes the Transition from Courtship to Copulation and Signals Female Acceptance in Drosophila melanogaster. Curr. Biol. 30, 3736–3748.e5 (2020).

24. Pavlou, H. J. et al. Neural circuitry coordinating male copulation. Elife 5, (2016).

25. Deutsch, D., et al. The neural basis for a persistent internal state in Drosophila females. eLife vol. 9 Preprint at 10.7554/elife.59502 (2020).

26. Chiu, H. et al. Cell type-specific contributions to a persistent aggressive internal state in female Drosophila. eLife 2023.06.07.543722 (2023) doi:10.1101/2023.06.07.543722.

27. Schretter, C. E. et al. Cell types and neuronal circuitry underlying female aggression in Drosophila. Elife 9, (2020).

28. Certel, S. J., Savella, M. G., Schlegel, D. C. F. & Kravitz, E. A. Modulation of Drosophila male behavioral choice. Proc. Natl. Acad. Sci. U. S. A. 104, 4706–4711 (2007).

29. Vrontou, E., Nilsen, S. P., Demir, E., Kravitz, E. A. & Dickson, B. J. fruitless regulates aggression and dominance in Drosophila. Nat. Neurosci. 9, 1469–1471 (2006).

30. Wang, F. et al. Neural circuitry linking mating and egg laying in Drosophila females. Nature 579, 101–105 (2020).

31. Vijayan, V. et al. A rise-to-threshold process for a relative-value decision. Nature 619, 563–571 (2023).

32. Rezával, C., Nojima, T., Neville, M. C., Lin, A. C. & Goodwin, S. F. Sexually dimorphic octopaminergic neurons modulate female postmating behaviors in Drosophila. Curr. Biol. 24, 725–730 (2014).

33. Vijayan, V. et al. An internal expectation guides Drosophila egg-laying decisions. Sci. Adv. 8, eabn3852 (2022).

34. Berg, S. et al. Sexual dimorphism in the complete connectome of the Drosophila male central nervous system. bioRxivorg (2025) doi:10.1101/2025.10.09.680999.

35. Dorkenwald, S. et al. Neuronal wiring diagram of an adult brain. Nature 634, 124–138 (2024).

36. Schlegel, P. et al. Whole-brain annotation and multi-connectome cell typing of Drosophila. Nature 634, 139–152 (2024).

37. Dorkenwald, S. et al. Neuronal wiring diagram of an adult brain. bioRxiv (2023) doi:10.1101/2023.06.27.546656.

38. Nojima, T. et al. A sex-specific switch between visual and olfactory inputs underlies adaptive sex differences in behavior. Curr. Biol. 31, 1175–1191.e6 (2021).

39. Chiang, A.-S. et al. Three-dimensional reconstruction of brain-wide wiring networks in Drosophila at single-cell resolution. Curr. Biol. 21, 1–11 (2011).

40. Wu, M. et al. Visual projection neurons in the Drosophila lobula link feature detection to distinct behavioral programs. Elife 5, e21022 (2016).

41. McKellar, C. E. et al. Threshold-Based Ordering of Sequential Actions during Drosophila Courtship. Curr. Biol. 29, 426–434.e6 (2019).

42. Stürner, T., et al. Comparative connectomics of Drosophila descending and ascending neurons. Nature (2025) doi:10.1038/s41586-025-08925-z.

43. Costa, M., Manton, J. D., Ostrovsky, A. D., Prohaska, S. & Gregory S X. NBLAST: Rapid, Sensitive Comparison of Neuronal Structure and Construction of Neuron Family Databases. Neuron vol. 91 293–311 Preprint at 10.1016/j.neuron.2016.06.012 (2016).

44. Matsliah, A., et al. Codex: connectome data explorer. Preprint at 10.13140/RG 2 (2023).

45. Sterne, G. R., Otsuna, H., Dickson, B. J. & Scott, K. Classification and genetic targeting of cell types in the primary taste and premotor center of the adult Drosophila brain. Elife 10, (2021).

46. Kimura, K.-I., Sato, C., Koganezawa, M. & Yamamoto, D. Drosophila ovipositor extension in mating behavior and egg deposition involves distinct sets of brain interneurons. PLoS One 10, e0126445 (2015).

47. Stürner, T. et al. Comparative connectomics of the descending and ascending neurons of the Drosophila nervous system: stereotypy and sexual dimorphism. bioRxivorg (2024) doi:10.1101/2024.06.04.596633.

48. Scheffer, L. K. et al. A connectome and analysis of the adult Drosophila central brain. Elife 9, (2020).

49. Matsliah, A. et al. Neuronal parts list and wiring diagram for a visual system. Nature 634, 166–180 (2024).

50. Laturney, M., Sterne, G. R. & Scott, K. Mating activates neuroendocrine pathways signaling hunger in Drosophila females. Elife 12, (2023).

51. Jung, Y. et al. Neurons that Function within an Integrator to Promote a Persistent Behavioral State in Drosophila. Neuron 105, 322–333.e5 (2020).

52. Court, R., et al. Virtual Fly Brain-An interactive atlas of the Drosophila nervous system. Front. Physiol. 14, 1076533 (2023).

53. Eckstein, N. et al. Neurotransmitter classification from electron microscopy images at synaptic sites in Drosophila melanogaster. Cell 187, 2574–2594.e23 (2024).

54. Imoto, K. et al. Neural-circuit basis of song preference learning in fruit flies. iScience 27, 110266 (2024).

55. Meissner, G. W., Luo, S. D., Dias, B. G., Texada, M. J. & Baker, B. S. Sex-specific regulation of Lgr3 in Drosophila neurons. Proc. Natl. Acad. Sci. U. S. A. 113, E1256–65 (2016).

56. Wang, F., Wang, K., Forknall, N., Parekh, R. & Dickson, B. J. Circuit and Behavioral Mechanisms of Sexual Rejection by Drosophila Females. Curr. Biol. 30, 3749–3760.e3 (2020).

57. Lee, G. et al. Spatial, temporal, and sexually dimorphic expression patterns of the fruitless gene in the Drosophila central nervous system. J. Neurobiol. 43, 404–426 (2000).

58. Stockinger, P., Kvitsiani, D., Rotkopf, S., Tirián, L. & Dickson, B. J. Neural circuitry that governs Drosophila male courtship behavior. Cell 121, 795–807 (2005).

59. Nern, A. et al. Connectome-driven neural inventory of a complete visual system. Nature (2025) doi:10.1038/s41586-025-08746-0.

60. Tanaka, R., Higuchi, T., Kohatsu, S., Sato, K. & Yamamoto, D. Optogenetic activation of the fruitless-labeled circuitry in Drosophila subobscura males induces mating motor acts. J. Neurosci. 37, 11662–11674 (2017).

61. Thistle, R., Cameron, P., Ghorayshi, A., Dennison, L. & Scott, K. Contact chemoreceptors mediate male-male repulsion and male-female attraction during Drosophila courtship. Cell 149, 1140–1151 (2012).

62. Toda, H., Zhao, X. & Dickson, B. J. The Drosophila female aphrodisiac pheromone activates ppk23(+) sensory neurons to elicit male courtship behavior. Cell Rep. 1, 599–607 (2012).

63. Datta, S. R. et al. The Drosophila pheromone cVA activates a sexually dimorphic neural circuit. Nature 452, 473–477 (2008).

64. Manoli, D. S. et al. Male-specific fruitless specifies the neural substrates of Drosophila courtship behaviour. Nature 436, 395–400 (2005).

65. Kamikouchi, A., Shimada, T. & Ito, K. Comprehensive classification of the auditory sensory projections in the brain of the fruit fly Drosophila melanogaster. J. Comp. Neurol. 499, 317–356 (2006).

66. Patella, P. & Wilson, R. I. Functional maps of mechanosensory features in the Drosophila brain. Curr. Biol. 28, 1189–1203.e5 (2018).

67. Baker, C. A. et al. Neural network organization for courtship-song feature detection in Drosophila. Curr. Biol. 32, 3317–3333.e7 (2022).

68. Vaughan, A. G., Zhou, C., Manoli, D. S. & Baker, B. S. Neural pathways for the detection and discrimination of conspecific song in D. melanogaster. Curr. Biol. 24, 1039–1049 (2014).

69. Zhou, C. et al. Central neural circuitry mediating courtship song perception in male Drosophila. eLife Sciences 4, e08477 (2015).

70. Jefferis, G. S. X. E. et al. Comprehensive maps of Drosophila higher olfactory centers: spatially segregated fruit and pheromone representation. Cell 128, 1187–1203 (2007).

71. Grosjean, Y. et al. An olfactory receptor for food-derived odours promotes male courtship in Drosophila. Nature 478, 236–240 (2011).

72. Kallman, B. R., Kim, H. & Scott, K. Excitation and inhibition onto central courtship neurons biases Drosophila mate choice. Elife 4, e11188 (2015).

73. Hindmarsh Sten, T., Li, R., Otopalik, A. & Ruta, V. Sexual arousal gates visual processing during Drosophila courtship. Nature 595, 549–553 (2021).

74. Dombrovski, M. et al. Molecular gradients shape synaptic specificity of a visuomotor transformation. Nature **644**, 453–462 (2025).

75. Cazalé-Debat, L. et al. Mating proximity blinds threat perception. Nature 634, 635–643 (2024).

76. Rings, A. & Goodwin, S. F. To court or not to court - a multimodal sensory decision in Drosophila males. Curr. Opin. Insect Sci. 35, 48–53 (2019).

77. Pacheco, D. A., Thiberge, S. Y., Pnevmatikakis, E. & Murthy, M. Auditory activity is diverse and widespread throughout the central brain of Drosophila. Nat. Neurosci. 24, 93–104 (2021).

78. Jeanne, J. M., Fişek, M. & Wilson, R. I. The Organization of Projections from Olfactory Glomeruli onto Higher-Order Neurons. Neuron 98, 1198–1213.e6 (2018).

79. Kohl, J., Huoviala, P. & Jefferis, G. S. Pheromone processing in Drosophila. Curr. Opin. Neurobiol. 34, 149–157 (2015).

80. Schlegel, P. et al. Information flow, cell types and stereotypy in a full olfactory connectome. Elife 10, (2021).

81. Namiki, S., Dickinson, M. H., Wong, A. M., Korff, W. & Card, G. M. The functional organization of descending sensory-motor pathways in Drosophila. Elife 7, (2018).

82. Chen, C.-L. et al. Ascending neurons convey behavioral state to integrative sensory and action selection brain regions. Nat. Neurosci. 26, 682–695 (2023).

83. Braun, J., Hurtak, F., Wang-Chen, S. & Ramdya, P. Descending networks transform command signals into population motor control. Nature 630, 686–694 (2024).

84. Bates, A. S., et al. Distributed control circuits across a brain-and-cord connectome. bioRxivorg (2025) doi:10.1101/2025.07.31.667571.

85. Zheng, Z. et al. A Complete Electron Microscopy Volume of the Brain of Adult Drosophila melanogaster. Cell 174, 730–743.e22 (2018).

86. Schretter, C. E. et al. Social state alters vision using three circuit mechanisms in Drosophila. Nature 637, 646–653 (2025).

87. Feng, K., Palfreyman, M. T., Häsemeyer, M., Talsma, A. & Dickson, B. J. Ascending SAG neurons control sexual receptivity of Drosophila females. Neuron 83, 135–148 (2014).

88. Coen, P. et al. Dynamic sensory cues shape song structure in Drosophila. Nature 507, 233–237 (2014).

89. Schlief, M. L. & Wilson, R. I. Olfactory processing and behavior downstream from highly selective receptor neurons. Nat. Neurosci. 10, 623–630 (2007).

90. Cheong, H. S. J. et al. Transforming descending input into motor output: An analysis of the Drosophila Male Adult Nerve Cord connectome. (2025) doi:10.7554/elife.96084.

91. Taisz, I. et al. Generating parallel representations of position and identity in the olfactory system. Cell 186, 2556–2573.e22 (2023).

92. Hindmarsh Sten, T., Li, R., Hollunder, F., Eleazer, S. & Ruta, V. Male-male interactions shape mate selection in Drosophila. Cell 188, 1486–1503.e25 (2025).

93. Weber-Langstaff, R. A., Srivastava, P., Kunin, A. B. & Gutierrez, G. J. The oviposition inhibitory neuron is a potential hub of multi-circuit integration in the Drosophila brain. eNeuro **12**, ENEURO.0123–25.2025 (2025).

94. Tootoonian, S., Coen, P., Kawai, R. & Murthy, M. Neural representations of courtship song in the Drosophila brain. J. Neurosci. 32, 787–798 (2012).

95. Tinbergen, N. The study of instinct Clarendon Press. Oxford 195, l (1951).

96. Pan, Y. & Baker, B. S. Genetic identification and separation of innate and experience-dependent courtship behaviors in Drosophila. Cell 156, 236–248 (2014).

97. Billeter, J.-C. & Goodwin, S. F. Characterization of Drosophila fruitless-gal4 transgenes reveals expression in male-specific fruitless neurons and innervation of male reproductive structures. J. Comp. Neurol. 475, 270–287 (2004).

98. Hoopfer, E. D., Jung, Y., Inagaki, H. K., Rubin, G. M. & Anderson, D. J. P1 interneurons promote a persistent internal state that enhances inter-male aggression in Drosophila. Elife 4, (2015).

99. Koganezawa, M., Kimura, K.-I. & Yamamoto, D. The Neural Circuitry that Functions as a Switch for Courtship versus Aggression in Drosophila Males. Curr. Biol. 26, 1395–1403 (2016).

100. Kohatsu, S. & Yamamoto, D. Visually induced initiation of Drosophila innate courtship-like following pursuit is mediated by central excitatory state. Nat. Commun. 6, 6457 (2015).

101. Ribeiro, I. M. A. et al. Visual Projection Neurons Mediating Directed Courtship in Drosophila. Cell 174, 607–621.e18 (2018).

102. Mabuchi, Y. et al. Visual feedback neurons fine-tune Drosophila male courtship via GABA-mediated inhibition. Curr. Biol. 33, 3896–3910.e7 (2023).

103. Lillvis, J. L. et al. Nested neural circuits generate distinct acoustic signals during Drosophila courtship. Curr. Biol. 34, 808–824.e6 (2024).

104. Shirangi, T. R., Stern, D. L. & Truman, J. W. Motor control of Drosophila courtship song. Cell Rep. 5, 678–686 (2013).

105. Otsuna, H., Ito, M. & Kawase, T. Color depth MIP mask search: a new tool to expedite Split-GAL4 creation. bioRxiv (2018) doi:10.1101/318006.

106. Bates, A. S. et al. The natverse, a versatile toolbox for combining and analysing neuroanatomical data. Elife 9, (2020).

107. Lin, A. et al. Network statistics of the whole-brain connectome of Drosophila. Nature 634, 153–165 (2024).

108. Schwartzman, G., Jourdan, B., García-Soriano, D. & Matsliah, A. NTAC: Neuronal type assignment from connectivity. Nat. Commun. 17, 1284 (2026).

109. Simpson, M., Srinivasan, V. & Thomo, A. Efficient computation of feedback arc set at web-scale. Proceedings VLDB Endowment 10, 133–144 (2016).

110. Meissner, G. W. et al. A split-GAL4 driver line resource for Drosophila neuron types. Elife 13, RP98405 (2025).

111. Pereira, T. D. et al. SLEAP: A deep learning system for multi-animal pose tracking. Nat. Methods 19, 486–495 (2022).

112. Cohen, N. A., Brenman, J. E., Snyder, S. H. & Bredt, D. S. Binding of the inward rectifier K+ channel Kir 2.3 to PSD-95 is regulated by protein kinase A phosphorylation. Neuron 17, 759–767 (1996).

113. Baines, R. A., Uhler, J. P., Thompson, A., Sweeney, S. T. & Bate, M. Altered electrical properties in Drosophila neurons developing without synaptic transmission. J. Neurosci. 21, 1523–1531 (2001).

114. Tao, L., Ayambem, D., Barranca, V. J. & Bhandawat, V. Neurons underlying aggression-like actions that are shared by both males and females in Drosophila. J. Neurosci. 44, e0142242024 (2024).

